# STAMP: Simultaneous Training and Model Pruning for Low Data Regimes in Medical Image Segmentation

**DOI:** 10.1101/2021.11.26.470124

**Authors:** Nicola K. Dinsdale, Mark Jenkinson, Ana I. L. Namburete

## Abstract

Acquisition of high quality manual annotations is vital for the development of segmentation algorithms. However, to create them we require a substantial amount of expert time and knowledge. Large numbers of labels are required to train convolutional neural networks due to the vast number of parameters that must be learned in the optimisation process. Here, we develop the *STAMP* algorithm to allow the *simultaneous* training and pruning of a UNet architecture for medical image segmentation with targeted channelwise dropout to make the network robust to the pruning. We demonstrate the technique across segmentation tasks and imaging modalities. It is then shown that, through online pruning, we are able to train networks to have much higher performance than the equivalent standard UNet models while reducing their size by more than 85% in terms of parameters. This has the potential to allow networks to be directly trained on datasets where very low numbers of labels are available.

## 1. Introduction

Semantic segmentation in medical imaging is vital for the understanding and monitoring of the progression of disease. For instance, the accurate segmentation of the hippocampus is essential for volumetric and morphological assessment, as hippocampal atrophy, observed through MRI, is one of the most validated biomarkers of Alzheimer’s disease (Flores et al., 2015). Manual segmentation, however, is time consuming and difficult. Several structures have ambiguous boundaries, making consistent delineation of the structures between raters hard to achieve. There is, therefore, a need for automated segmentation methods, capable of reliably providing accurate segmentations of the structures of interest.

Deep learning-based methods have become state-of-the-art for medical image segmentation, with most methods being based on the UNet architecture (Ronneberger et al., 2015; Cicek et al., 2016; Milletari et al., 2016). Methods either using UNets directly, or highly inspired by the UNet, have been applied to the spectrum of medical imaging segmentation tasks across a range of modalities: for example, in MRI (Guha Roy et al., 2018; Balakrishnan et al., 2019; Isensee et al., 2021; Dinsdale et al., 2019), in CT (Isensee et al., 2021; Schlemper et al., 2019), in X-Ray (Isensee et al., 2021; Yahyatabar et al., 2020) and in ultrasound (Schlemper et al., 2019; Vaze et al., 2020).

A major limitation of these methods, and with deep learning-based methods in general, is the large amount of labelled data required to train the models. In medical imaging, manual delineation by domain experts is considered to be the ‘gold standard’ for labels. Therefore, to produce a dataset large enough to train a network for segmentation is expensive, requiring large amounts of expert time to curate, and expert domain knowledge. Furthermore, the labels required often do not form part of standard clinical practice, and therefore have to be produced specifically for the network training. Thus, there is a need to develop methods that can work in data domains where low numbers of labelled data points are available.

One reason why large training dataset sizes are needed is that the models contain large numbers of parameters – many more than the number of training examples available. Many methods have been proposed to help to train models in low data regimes, with the most frequently used approaches being model pretraining or transfer learning (Raghu et al., 2019), and data augmentation (Nalepa et al., 2019). Model pretraining uses a related large dataset to initialise the model weights, allowing the optimisation for the target dataset/task to begin from a more informed place. Data augmentation applies random transformations to the data during training, increasing the size and variation of the dataset seen, thus artificially creating more data points to train the model. We propose to investigate the utility of *model pruning*, reducing the number of parameters in the model and so the model complexity, and thus potentially reducing the number of data points required to train the model.

Model pruning for neural networks was first explored in (Cun et al., 1990), where they introduced the standard pruning framework. A model is trained until it performs acceptably well on a given task and then the weights with small saliency – that is, the weights that have the least effect on the error – are removed and then this pruned network is finetuned. The process is repeated iteratively until a model of the desired size or performance is obtained. This general format is replicated across the literature; however, the details of each stage vary between the study and the applications.

First, calculation of which weights have the least effect on the model error is computationally highly expensive (Molchanov et al., 2016) and so an approximation is desirable. Various metrics have been proposed to approximate this saliency, including the second derivative of the gradient (Cun et al., 1990), the magnitude of the weights or activations (Han et al., 2015; Frankle and Carbin, 2019; Hu et al., 2016; Li et al., 2017), and the product of the activation and gradients (Molchanov et al., 2016).

Methods also differ as to the mechanism of the pruning. Some methods prune individual model weights or connections (*unstructured pruning*) (Cun et al., 1990; Frankle and Carbin, 2019; Srinivas and Venkatesh Babu, 2015) whereas others prune whole convolutional filters (*structured pruning*) (Molchanov et al., 2016; Li et al., 2017). Unstructured pruning results in networks that are sparse and it is hard to accelerate these models without specialised libraries, making them difficult to apply using general-purpose hardware (Molchanov et al., 2016). Thus, this work will focus on structured pruning: removing whole convolutional filters. We will also follow the work of (Li et al., 2017) and remove whole filters from the model during training time, such that the model becomes progressively smaller.

The methods across the literature prune the network after training to convergence, then fine-tune the resulting model. This, however, requires the model to be trained to convergence – or at least to good performance by some metric (Cun et al., 1990) – before we are able to prune the network. This leads to long training times and the methods potentially not being applicable if one cannot train the full-sized network to an acceptable performance in the first place. Few methods attempt to prune the model during training (Bellec et al., 2017; Shen et al., 2020). Other methods exist that have been applied to medical imaging, which aim to create networks with inherently reduced numbers of parameters, such as model distillation (Vaze et al., 2020; Hinton et al., 2014) and the use of separable convolutions (Vaze et al., 2020; Howard et al., 2017).

The obvious question, given that it is often possible to prune networks to less than 50% of the original size of the model, is whether the pruning process can be avoided altogether by simply training smaller models directly. It has, however, been shown by several works in the literature that it is not normally possible to train the smaller pruned networks from random initialisation and achieve the same performance (Han et al., 2015; Li et al., 2017). The Lottery ticket hypothesis (Frankle and Carbin, 2019) demonstrates that there exists a subnetwork of the original network that can be trained to the same level of task performance, given that the starting point is a dense network reduced using unstructured pruning, if a suitable initialisation could be found. The subnetwork is then a pruned network, taken in isolation, with the rest of the network removed. These networks are very sensitive to initialisation and smaller networks may be achievable through maintaining the weights of the pruned networks, due to coadapted layers from earlier in the pruning process maintaining the complex relationships that were learned. The sensitivity to the initialisation when pruning convolutional filters as opposed to individual parameters is demonstrated in (Cohen et al., 2016).

Existing pruning methods have been validated in high data regimes, primarily on large computer vision classification datasets such as MNIST (Lecun et al., 1998) – (Cun et al., 1990; Frankle and Carbin, 2019; Hu et al., 2016) – CIFAR10 (Krizhevsky et al., 2009) – (Frankle and Carbin, 2019; Li et al., 2017) – and ImageNet (Deng et al., 2009) – (Han et al., 2015; Hu et al., 2016). Datasets of this size do not exist in medical imaging, where manual annotations are expensive and time consuming to acquire, and datasets are more commonly of the order of hundreds of labelled examples. We therefore propose to explore using model pruning for training in low data regimes: specifically, our algorithm simultaneously trains and prunes a UNet architecture. The utility of pruning in very low data regimes is explored here for a range of segmentation tasks and datasets for medical imaging.

Our contributions, therefore, are as follow:

- We propose an algorithm to *simultaneously* train and prune a UNet. It is, to the best of our knowledge, the first exploration of pruning applied to a UNet for medical image segmentation, removing whole convolutional filters through the pruning process. We demonstrate our technique on several open access datasets and segmentation tasks, across imaging modalities.
- A channelwise targeted dropout is developed, based on that proposed in (Gomez et al., 2018), but adapted to work channelwise and without requiring availability of information about the filters that are most likely to be removed before training. This is similar to the method proposed in (Hou and Wang, 2019), where channelwise weighted dropout is used to regularise the training of a model, but the normalised filter magnitudes calculated for pruning are used to modulate the dropout. It is demonstrated that this improves the performance and stability of the models during training, by making them more robust to the pruning process.
- Finally, we explore model pruning for very low data regimes, and we show that, through training and pruning the model simultaneously, it is possible to achieve significantly higher performance than can be obtained by directly training UNet models of the equivalent (final) size.

## 2. Methods

Our aim is to train a deep neural network to segment regions of interest from input images while limiting the degree of overfitting, by pruning the model during training. We hypothesize that the trained model should require *less* training data points to produce good quality segmentations, due to the reduction in parameters that must be optimised. As this work uses medical images, where labels are expensive to acquire, it must function in a low data regime. Our discussion will largely follow the notation introduced in (Molchanov et al., 2016).

To allow the network to be trained and pruned simultaneously, consider the scenario where there is access to a training dataset 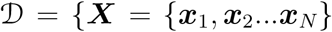, ***Y*** = {***y***_1_, ***y***_2_…***y***_*N*_}} where 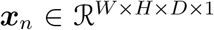 represents an input image and 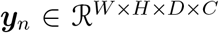 is its corresponding target segmentation, and where *C* corresponds to the number of classes in the target segmentation. A network is trained to predict the target segmentations ***y***_*n*_ from the input images ***x***_*n*_ and the network is parametrised by 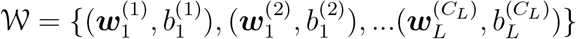 where 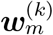 corresponds to the weight kernel for the *k^th^* filter of the *m^th^* layer and 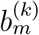 is the corresponding bias that will be incorporated into discussion of weights from here on. *C_L_* is the number of channels in layer *L*. The weights are first randomly initialised and are optimised during training by minimising a loss function 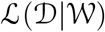 that aims to optimise the performance on the segmentation task and can be chosen completely independently of this pruning process.

During the pruning process, the aim is to refine the parameters of the network W to a smaller subset of the parameters 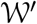 such that 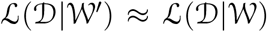 (Molchanov et al., 2016). To achieve this, we use the observation that the parameters of the smallest magnitude generally have the least impact on the final segmentation produced by the network, and can be removed with the least impact on model performance (Cun et al., 1990).

### 2.1. Training Procedure

The overall training procedure is shown in Fig. 1. The model weights are initialised with Xavier initialization (Glorot and Bengio, 2010) and the model is trained for a single epoch before pruning begins, so that the filters contain some information about the task and not just the initialised weights.

**Figure 1:**
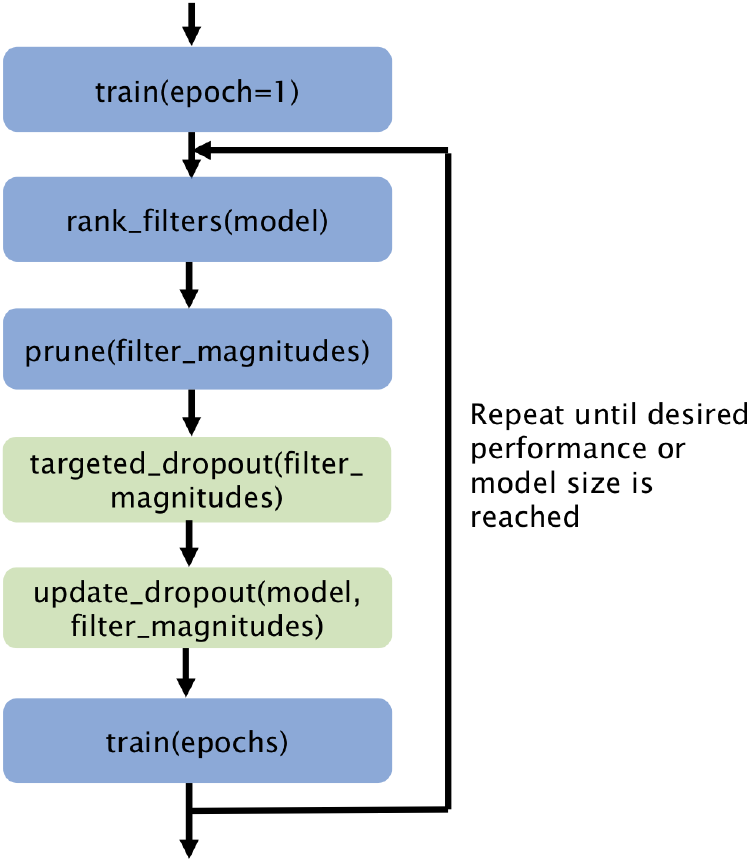
Overall *STAMP*+ training procedure for training and pruning a model simultaneously. The targeted dropout algorithm is presented in Algorithm 1. Steps in green are removed in ablation experiments.

Our pruning strategy removes whole convolutional kernels as shown in Fig. 2, meaning that we do not end up with a sparse representation and are able to use standard libraries and hardware. When pruning whole convolutional filters, we remove the filter from the kernel matrix ***w***_*i*_, the corresponding bias, and the filter from the weight kernel matrix ***w***_*i*+1_, as in (Li et al., 2017). To decide which filters to remove, we consider the feature activation maps, denoted by 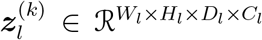 where *l* ∈ {1,…, *L*}, is the current layer, and *C_l_* is the number of channels at layer depth *l*. The activation maps are related to the kernel weights by:

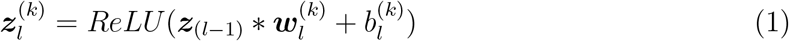

where ReLU is the activation function used throughout our network, other than for the final convolution. The use of ReLU means that our network contains a substantial number of zero-valued parameters, creating redundancy that can be pruned.

**Figure 2:**
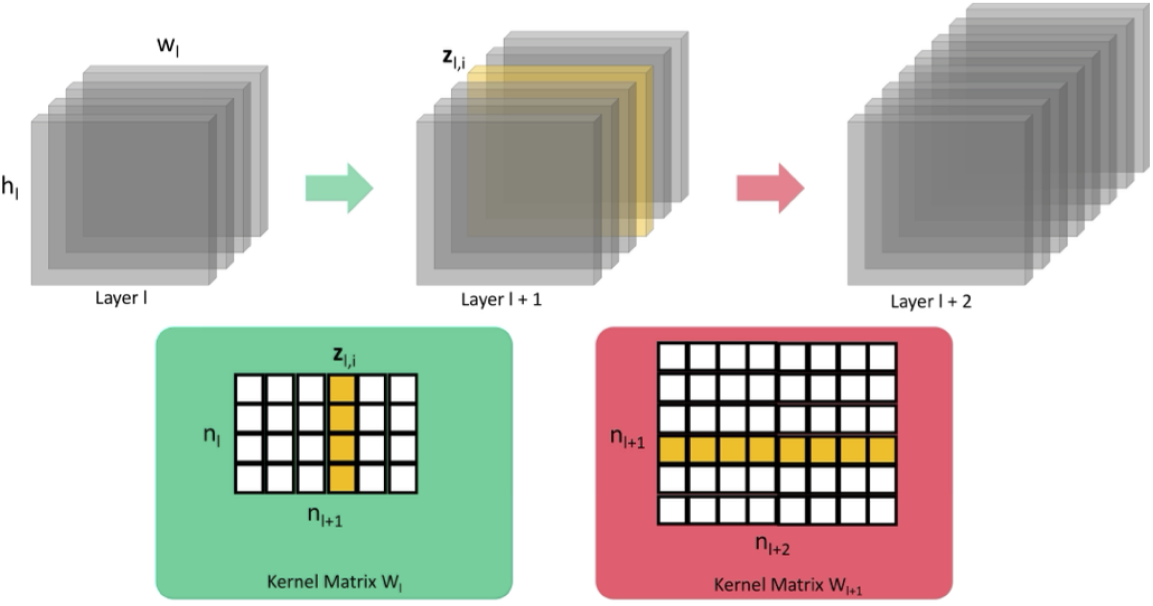
Whole convolutional filters are pruned, based on the magnitude of the corresponding feature maps ***z***_*l,i*_ (the convolutional kernel to be removed is shown in yellow). This requires the weights to be pruned from the kernel matrix ***w***_*l*_ and ***w***_*l*+1_ (shown in yellow in the kernel matrices). The corresponding biases are also pruned but are not shown.

We use the observation that weights with the smallest impact on the final prediction are those with the smallest magnitudes (Cun et al., 1990), and so the weights, 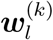, and biases 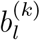, corresponding to the smallest magnitude filter activations, 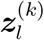, are pruned from the network. To assess the overall magnitude of the filters, the *L*2 norm (Han et al., 2015), averaged across all training data points (calculated from an additional forward pass), is considered:

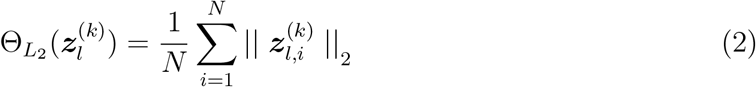

The *L*2 norm is used as it is computationally simple and provides stable performance, but other metrics could be used to evaluate the magnitude of the filters. This was explored in the supplementary material, and it was found that *STAMP* was robust to this choice.

These values must then be normalised across the layer depth, because the values are at different scales at different depths (Molchanov et al., 2016). Therefore, a simple *L*2 normalisation across the values at each layer is employed.

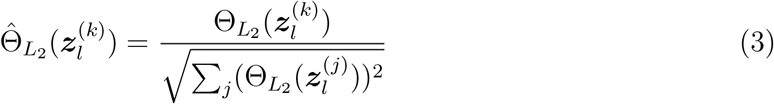

where *j* iterates over all the filter kernels in layer *l*.

To prune the filter, it is removed entirely from the model architecture, rather than just set to zero, and so the model architecture reduces in the number of parameters as the training progresses (Li et al., 2017). Practically, this is achieved by creating a smaller model with the selected filter removed and then reloading all of the weights apart from those corresponding to the pruned filter.

### 2.2. Adaptive Channelwise Targeted Dropout

To make the model more robust to the pruning, *adaptive channelwise targeted dropout*, based on (Gomez et al., 2018) and (Hou and Wang, 2019), is introduced. The goal is to create a dropout scheme where the convolutional kernels which are most likely to be removed by the pruning procedure are the most likely to be dropped out during training.

#### Algorithm 1

Adaptive Targeted Dropout Algorithm. The *index* function returns the index (or position) of the Θ for the *k^th^* filter at layer *l* within the full sorted list of Θ values across the network. From this, the average index at each layer depth is calculated and used for determining the dropout probability (***p***) for each layer. The output ***p*** is a vector of probabilities. [] represents a list of elements, such that they remain together during the *sort_descending* process.

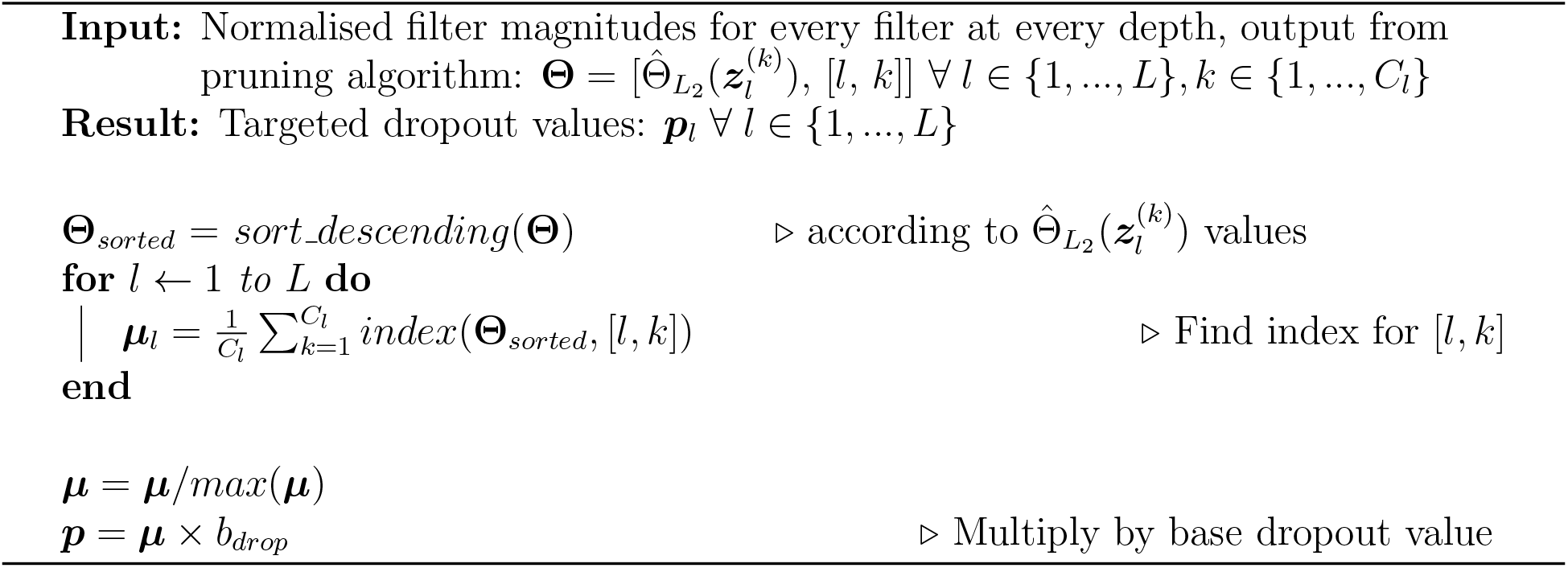

Our approach builds upon two previous works. In (Gomez et al., 2018) parameterwise dropout is applied to units which are *a priori* believed to be the least useful, thus encouraging the network to learn a representation that is more robust to *post-hoc* sparsification. In (Hou and Wang, 2019) weighted channelwise dropout is applied during training to regularise the model training. Global Average Pooling is used to determine the importance of individual filters, and a different dropout value is applied to each filter. Here, we apply the same dropout value to all of the filters at a given layer depth such that the information carried by the pruned filters is removed, and not redistributed to other filters at that depth, otherwise the model will continue to overfit to the training data. Like (Hou and Wang, 2019) the dropout is applied channelwise, rather than parameter-wise (Gomez et al., 2018), so that the network is prepared for the removal of whole convolutional filters; thus, channelwise or *spatial* dropout (Tompson et al., 2015) is added to our model. We do not know which filters are the most likely to be pruned *a priori* and, due to not having a pretrained model which is then pruned, the filter magnitudes cannot be used as a basis to predict this before the process begins. Therefore, unlike in (Gomez et al., 2018), the targeted dropout values cannot be based on a known probability distribution.

Thus, we develop our targeted dropout scheme as follows and as shown in Algorithm 1:

- The dropout values are changed adaptively during training, based on the calculated filter magnitudes, 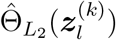.
- After the pruning has been completed, the new values of 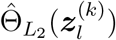 are calculated. These normalised magnitudes are then ordered.
- For each layer depth in the network, the average indices, *μ_l_* of the ordered normalised filter magnitudes are calculated (i.e. the average position in the sorted list of the filters at a given depth). This was based on empirical findings that the discrete values from the filter indices were better at encoding the relative likelihoods of the filters being pruned from the model.
- Each *μ_l_* is then normalised by the maximum value, such that the values are between 0 and 1. These values are then used to modulate *b_drop_* such that the new dropout probability for layer *l* is given by 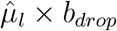.

When the new dropout values, ***p***, are calculated, the model architecture is updated with the new spatial dropout values, with the same value being applied to all filters in a layer.

### 2.3. Model Training

After pruning and updating the targeted dropout values, the model is then trained using standard backpropagation. The number of epochs between filter prunings, described as *recovery epochs*, is a hyperparameter that needs to be decided, and its effect on the segmentation performance will be explored.

In the experiments, the network is trained until it cannot be pruned any more, to allow exploration of the model’s performance, even as the model becomes very small. A model is considered to have reached the pruning limit when only a single filter remains at each depth and so to prune further would be to break the model. In practice, the model could be trained with standard early stopping, to save the best performing model.

### 2.4. Network Architecture

Across the experiments, a standard 2D or 3D UNet architecture is explored (Ronneberger et al., 2015; Cicek et al., 2016) (Fig. 3) with two blocks of convolutional layers at each depth, each with a batch normalisation layer (Ioffe and Szegedy, 2015). The UNet architecture is considered as it is widely used throughout medical image segmentation, and many winning architectures in segmentation challenges are based on it. Only the basic UNet is considered, but it is expected that the findings should generalise well to UNet-derived (encoder-decoder) architectures.

**Figure 3:**
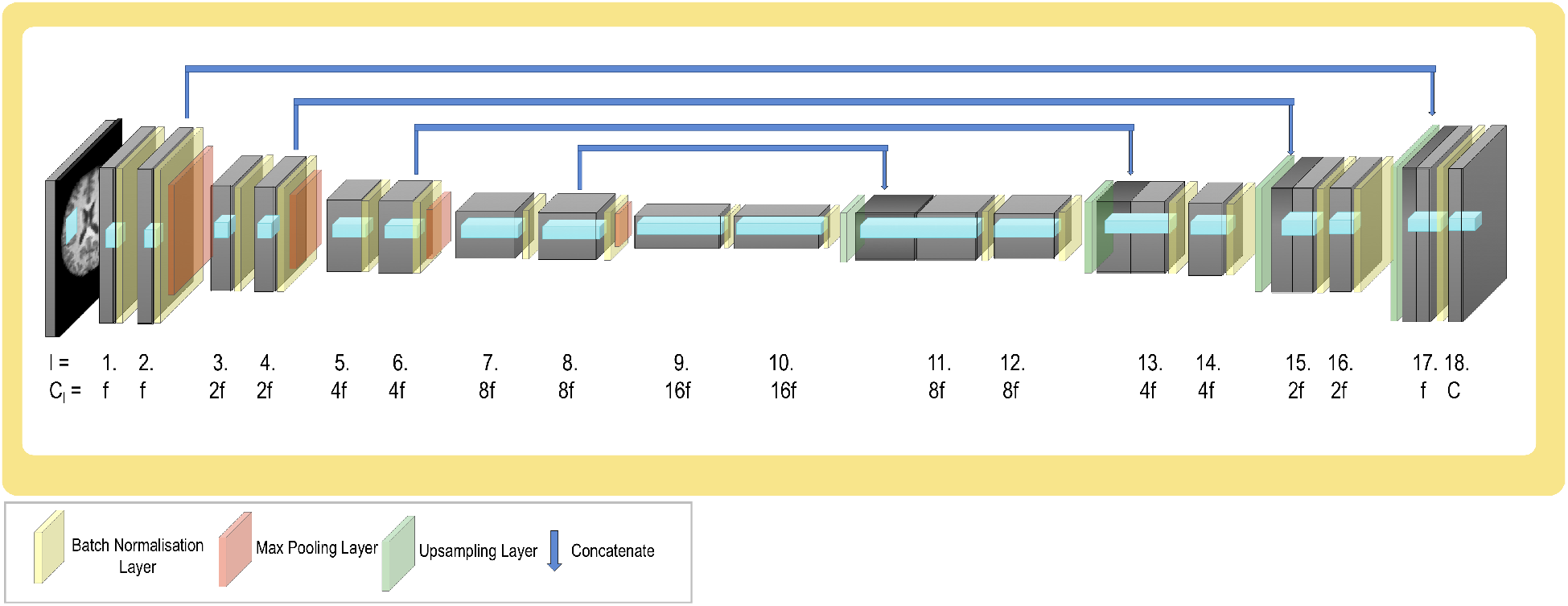
UNet model architecture: it follows the standard pattern of halving resolution and doubling filters at each depth. *l* corresponds to the layer depth, *C_l_* is the number of channels in that layer and *C* is the number of classes in the output segmentation. *f* is the initial number of filters, and is varied across experiments, but is 4 unless otherwise stated.

ReLU activations are used throughout, apart from the final layer, where a sigmoid activation function is used to create the segmentations. ReLU is very commonly used in network architectures, and, when we are considering pruning, comes with the added advantage that it encourages the network parameters to become sparse, so filter values on average become lower. Spatial dropout is applied to each convolutional block during training, with the values determined by the Targeted Dropout algorithm explained in Algorithm 1.

The final layer (*l* = 18) contains the same number of filters as output classes *C* and so clearly cannot be pruned (in all cases the background class is considered as an additional class). Therefore, only filters in layers 1 – 17 will be considered as candidates to be pruned and therefore for Eq. (1) is evaluated over *l* ∈ {1,…, *L*} where *L* = 17 and the final layer is not considered.

The architecture follows the conventional pattern of halving resolution and doubling features at each depth. The value of *f*, which determines the number of filters in each layer, is varied in the experiments, allowing exploration of the effect of the initial model size on the pruning process and the model’s performance. The maximum size of the network architecture explored was determined by the available GPU memory. A shallower UNet was also considered, the results of which can be seen in the supplementary material.

### 2.5. Implementation Details

#### 2.5.1. Hyperparameters and Baselines

The hyperparameters were set as follows, unless otherwise stated:

- Recovery Epochs = 5
- *b_drop_* =0.1
- Initial number of filters f = 4
- Batch size = 16
- Learning rate = 0.01 with Adam Optimiser

and the effect of the hyperparameters introduced by our work, recovery epochs and *b_drop_*, is explored in Section 3.4.

For the various experiments, two variations of the proposed method and two benchmarks were considered, which will be referred to as follows:

- *STAMP* - The proposed method, training and pruning the network simultaneously, without the green blocks shown in Fig. 1, the procedure therefore consisting solely of calculating the 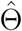 values and pruning the smallest filter;
- *STAMP*+ Targeted Dropout (*STAMP*+) - The proposed method, including the targeted dropout – all of the blocks in Fig. 1. These first two methods allow exploration of the impact of the pruning and the targeted dropout separately;
- *Standard UNet* - The first baseline, in which the UNet model is trained to convergence, without pruning, with the same hyperparameters and a patience of 25 epochs, representing the standard approach;
- *PruneFinetune* - The second baseline, in which a UNet that is trained to convergence, without pruning, is used for initialisation, and then the model is pruned using the same framework as for *STAMP*+. A patience of 10 epochs was used during the fine-tuning stages. This is the classic pruning regime, allowing exploration of the advantage of *STAMP*+, when pruning and training simultaneously.

The *PruneFinetune* baseline was chosen to be a fair comparison to standard pruning methods. Given that none of the existing methods were developed for UNet-style architectures and segmentation tasks, nor for low data regimes, implementing the methods as presented in prior work was not appropriate. Thus, we retained the pruning metric and pruning quantity from *STAMP+*, and thus these were combined with the standard procedure of taking the pretrained model, pruning the filter and then fine-tuning to convergence. In this way we were best able to compare any advantage of simultaneously training and pruning, and the results were unlikely to be due to other design decisions.

The same data splits were used for evaluating all methods. Five-fold cross-validation (the training data was split 80% training, 20% validation) was used for all experiments and all reported values used a further held-out test set for each dataset. No augmentation was applied to the data at any stage. We reported Dice scores throughout, but it can be found in the supplementary material that the results were consistent across other common evaluation metrics. All reported statistics were paired t-tests. All graphs show the mean results and interquartile range.

When a single value was reported for the performance of the pruning methods, the best model was chosen on the basis of results obtained from the validation data. All of the results were then reported for the held-out testing data, using the selected model.

The code was implemented using Python 3.5.2 and PyTorch 1.0.1.post2 (Paszke et al., 2019). All models were trained using a V100 GPU. The code is available at github.com/nkdinsdale/STAMP.

#### 2.5.2. Datasets

Seven medical imaging datasets were chosen for these experiments, spanning a range of tasks and imaging modalities. Basic details are listed in Table 1.

**Table 1:** Details of the datasets used in pruning experiments.

| Dataset | Modality | 2D/3D | Task | Image Size | Training/<br>Testing Subjects |
| --- | --- | --- | --- | --- | --- |
| HarP (Frisoni and Jack, 2015) | T1 MRI | 3D | Hippocampal Segmentation | $64 \times 64 \times 64$ | 200/70 |
| Medical Decathlon - Cardiac (Tobon-Gomez et al., 2015) | bSSFP MRI | 2D | Right Atrium | $320 \times 320$ | 15/5 (split into slices) |
| Medical Decathlon - Spleen (Simpson et al., 2014) | CT | 2D | Whole Spleen | $512 \times 512$ | 28/10 (split into slices) |
| Medical Decathlon - Prostate (Litjens et al., 2012) | Multimodal MR (T2 & ADC) | 2D | Whole Prostate | $320 \times 320$ | 28/10 (split into slices) |
| INTERGROWTH-21<sup>st</sup> Fetal Ultrasound (Papageorghiou et al., 2014) | Ultrasound | 3D | Fetal Brain | $160 \times 160 \times 160$ | 822 / 274 |
| IXI - HH | T1 MRI | 3D | Brain Extraction | $128 \times 128 \times 128$ | 166/19 |
| IXI - Guys | T1 MRI | 3D | Subcortical Segmentation | $128 \times 128 \times 128$ | 257/65 |

##### HarP - Brain

T1 MRI images, centred around the hippocampus, with 200 3D images for training (left and right hippocampus separately) and 70 for testing. The task was the segmentation of the hippocampus using manual labels, such that *C* = 2 (hippocampus and background). The dataset was originally reported in (Frisoni and Jack, 2015).

##### Medical Decathlon - Cardiac

MR images (bSSFP) covering the entire heart split into slices (varying numbers per image, 1771 in total for training), with 15 subjects for training and 5 for testing. The task was the segmentation of the right atrium, such that *C* = 2 (right atrium and background). The dataset was originally reported in (Tobon-Gomez et al., 2015).

##### Medical Decathlon - Spleen

CT dataset split into slices (varying numbers per image, 2691 in total for training), with 28 subjects for training and 10 subjects for testing. The task was the segmentation of the spleen, using automated segmentations that were manually corrected, such that *C* = 2 (spleen and background). The dataset was originally reported in (Simpson et al., 2014).

##### Medical Decathlon - Prostate

Multimodal MRI dataset (T2 and ADC) split into slices (varying numbers per image, 1648 in total for training), with 28 subjects for training and 10 subjects for testing. The task was the segmentation of two adjacent regions of the prostate, such that *C* = 3 (two prostate regions and background). The dataset was originally reported in (Litjens et al., 2012).

##### INTERGROWTH-21^*st*^ - Brain

Fetal ultrasound dataset, with 822 ultrasound 3D images for training and 274 for testing. The task was the segmentation of the fetal brain, using brain masks created by manually aligning the age matched fetal MRI atlas (Gholipour et al., 2017) to each US image, such that *C* =2 (fetal brain and background). The dataset was originally reported in (Papageorghiou et al., 2014).

##### IXI^1^ - HH

3T T1 MRI dataset, preprocessed using FSL Anat^2^, with 3D images of 166 subjects for training and 19 subjects for testing. The task was segmentation of the brain, with the labels being automatically generated by the FSL Anat pipeline by non-linearly registering an atlas to the target image, such that *C* = 2 (foreground and background).

##### IXI - Guys

1.5T T1 MRI dataset, preprocessed using FSL Anat, with 3D images of 257 subjects for training and 65 for testing. The task was segmentation of three deep matter subcortical structures: caudate, putamen and thalamus, with the labels automatically generated using FSL FIRST (Patenaude et al., 2011) run on the bias-field corrected and registered images output from the FSL Anat pipeline, such that *C* = 4 (caudate, putamen and thalamus, and background).

## 3. Results

The following section will explore first the low data regime results from across the datasets (Section 3.1), including an ablation study (Section 3.2). Then we present a methods comparison (Section 3.3), comparing *STAMP+* to *PruneFinetune*, smaller initial models and training the intermediate pruned models from scratch. Finally, we explore the effect of the introduced hyperparameters (Section 3.4): the number of recovery epochs and the *b_drop_* value.

### 3.1. Low Data Training

We first explored the performance of *STAMP+* in low data regimes, across a range of tasks and imaging modalities. For each modality we compared to the *Standard UNet*, and showed example segmentations for the two methods on the full dataset. Figure 4 shows the results on the HarP data, where it can be seen that *STAMP+* outperforms the *Standard UNet* across the range of training dataset sizes considered, with the performance of both methods improving as more training data is available, as would be expected. *STAMP+* also produced more consistent segmentation results, especially when only small amounts of training data were available. Figure 4 also shows the best, worst, and average Dice scores for 200 data points. Considering the poor and average segmentation performance, it is evident that *STAMP+* is better able to recover the fine features of the segmentation.

**Figure 4:**
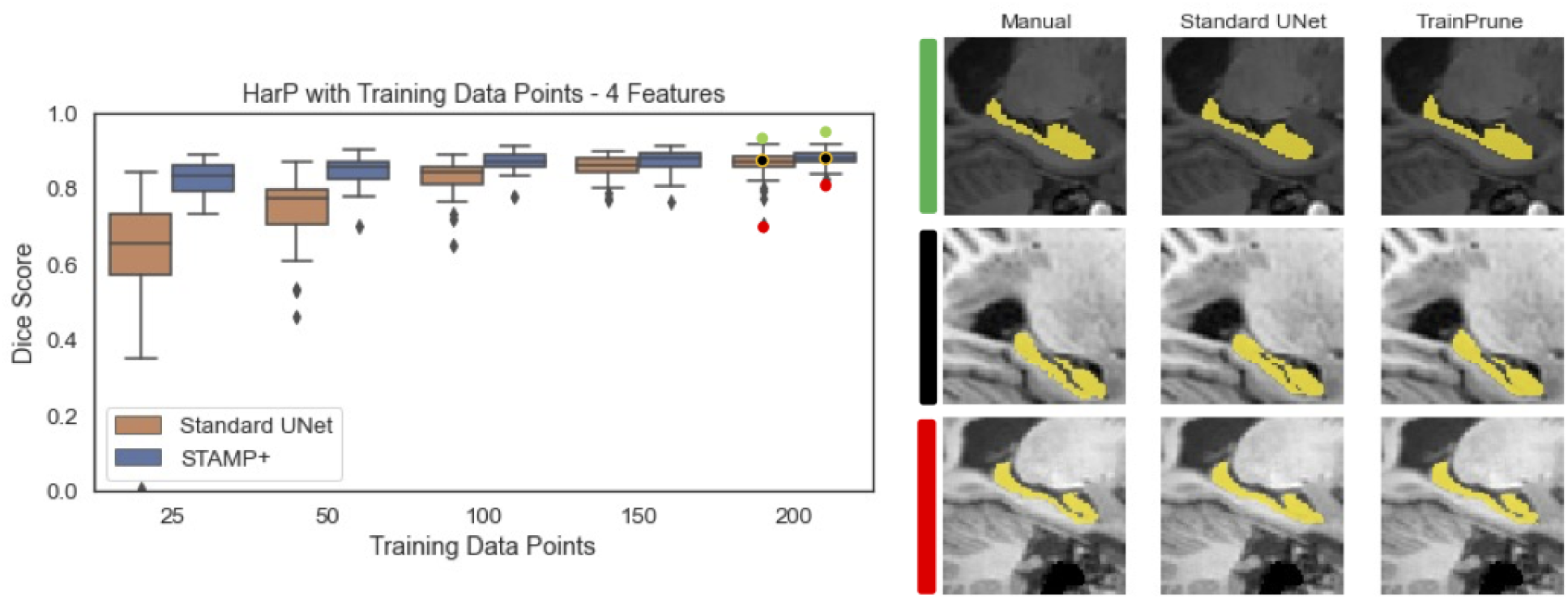
Dice scores for the HarP hippocampal segmentation for increasing amounts of training subjects, comparing *STAMP+* and the Standard UNet. Representative segmentations are also shown, for the best, worst, and average Dice scores.

Figure 5 shows the Dice scores on the HarP data plotted against the number of parameters remaining in the model. The mean value across the test set is shown as the solid line, and the shaded region indicates the interquartile range. The results were plotted against the remaining parameters rather than the pruning iteration. Different pruning iterations removed different numbers of parameters, depending on the location in the network from which the filter was removed: for instance, if it also led to filters being removed across the skip connection. It is evident that *STAMP+* was able to create models which outperformed the *Standard UNet* baselines, through training while removing model parameters. The improvement was greater when working in low data regimes, although the performance was less stable between iterations. Across the box plot results, we reported the Dice scores on the testing data from the model selected as best-performing based on the validation data. This pattern was observed across the datasets explored. Through plotting the Dice score versus parameters remaining we are able to compare the stability of training *STAMP+*, where it can be seen, through comparing the dice score between pruning iterations, that the training of *STAMP+* was more stable with more training data. We can also see that more parameters have to be removed before *STAMP+* reaches a high dice score on the testing data. Finally, we can see that the model maintains the high dice performance until the model becomes very small in terms of parameters, both with 200 and 50 data points for training.

**Figure 5:**
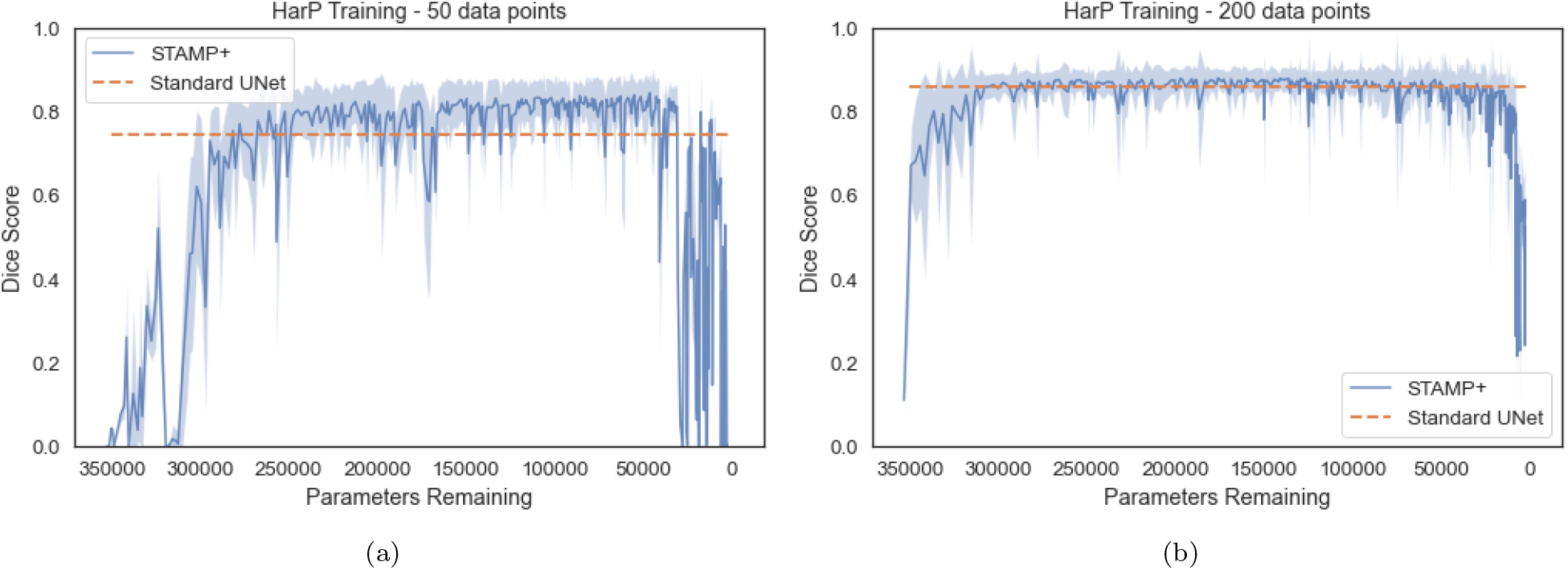
*STAMP+* segmentation results on the HarP data for both 50 data points for training and 200, with dice score plotted against the number of parameters remaining in the network architecture. As the network is pruned, the number of parameters reduces, and so the x-axis is inverted. The mean dice score is shown, with interquartile range. The mean dice score from the *Standard UNet* is shown for comparison. Note that *STAMP+* begins from random initialisation and so the performance is initially poor. It is evident that the performance improvement is greatest with a low number of data points but the performance is more stable between iterations with more training data.

We also considered five further datasets with the results shown in Fig. 6. It can clearly be seen that across the segmentation tasks and imaging modalities, *STAMP+* outperformed the *Standard UNet* when there were low numbers of training subjects, showing the power of the pruning method for working in low data regimes. It was most striking for the lowest number of subjects, for instance for the cardiac dataset (Fig. 6(a)): the *Standard UNet* was entirely unable to segment when training with 50 and 100 slices, whereas *STAMP+* was able to complete the segmentation to a similar standard as the *Standard UNet* when presented with more than double the number of samples.

**Figure 6:**
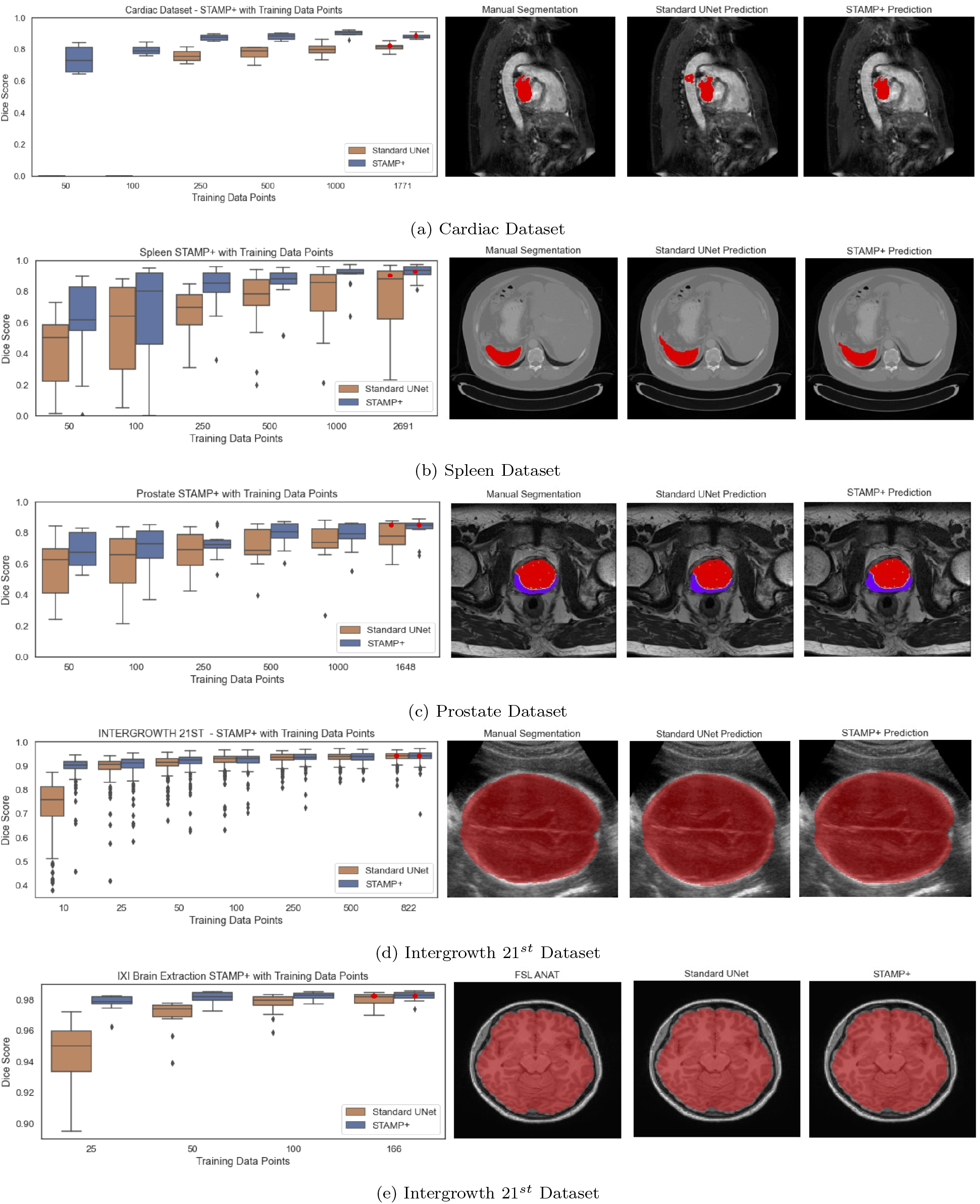
Dice scores for training low numbers of subjects comparing *STAMP+* and the Standard UNet for each dataset. Note that the Standard UNet was unable to complete the segmentation for the two lowest numbers of data points with the cardiac dataset. The qualitative example shown represents the median performance for *STAMP+* with the whole dataset for training.

### 3.2. Ablation Study: IXI Subcortical Segmentation

The final modality and task considered was segmenting three deep grey matter structures from T1 MRI: the caudate, putamen and thalamus. Without data augmentation, it was not possible to train a *Standard UNet* to successfully segment all three regions; however, it was possible to train the *STAMP+* method to segment all three regions. Therefore, an ablation study was performed to explore the effect of the pruning and the targeted dropout on the performance of the model. The following methods were tested: *Standard UNet*, *Standard UNet* with channelwise dropout, *Standard UNet* with targeted dropout, *STAMP*, *STAMP* with channelwise dropout (*STAMP+D*), and *STAMP+*. For the models with spatial dropout, the dropout probability was set to the average value of the targeted dropout values. Even for the relatively simple task of brain segmentation for the ultrasound data, there was significant improvement when using *STAMP+* for very low numbers of data points (ie. 10 data points).

First, the training distributions for the three pruning studies can be seen in Fig. 7. This shows that the distributions of the remaining filters throughout the pruning process as the models were pruned were very similar. This demonstrated that the targeted dropout was not driving the pruning process, with all methods showing the same pattern, and so we conclude that the pruning was being driven by the data. The improvement seen through the addition of the targeted dropout can be attributed to its ability to improve the model’s robustness to filter removal.

**Figure 7:**
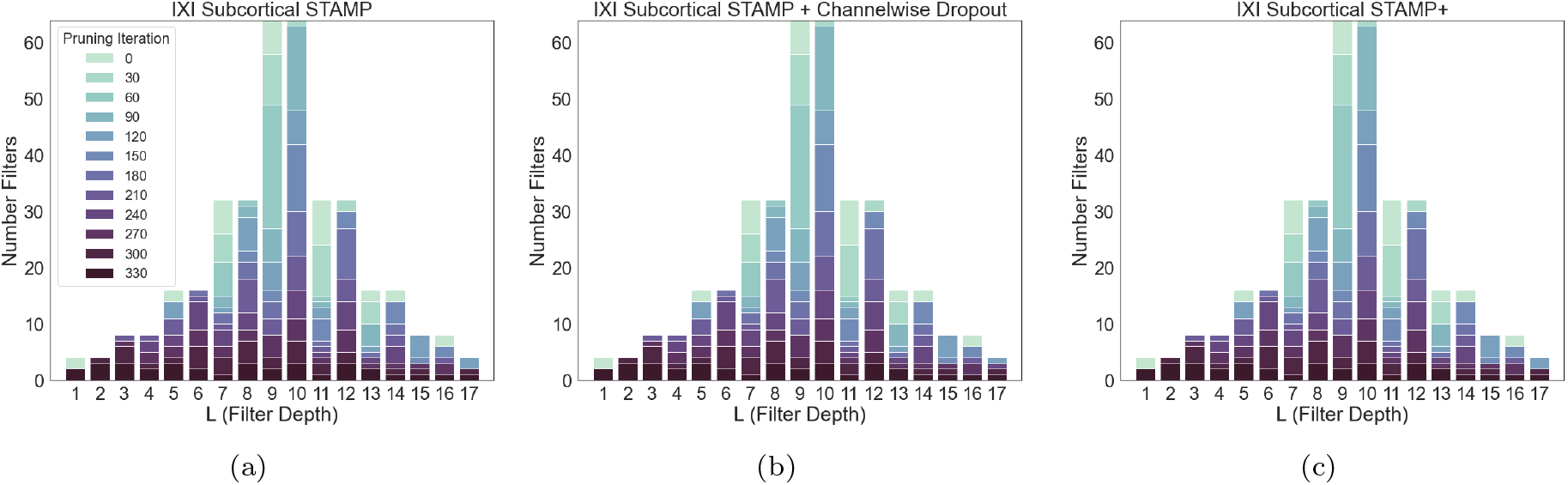
(a) *STAMP*, (b) *STAMP* + *Channelwise dropout out* (c) *STAMP+* for subcortical segmentation on the IXI data. The darker the plot, the longer the filters at that depth were kept in the model (l (filter depth) corresponds to the values shown on figure 3). Plots (a) to (c), represent increasing complexity in the applied regularisation, through dropout, to the model. It can be seen that all the methods led to a very similar pattern of pruning of the filters. Therefore, the adaptive dropout did not drive which filters are pruned, merely made the model more robust to being pruned, showing that the pruning pattern is a function of the training data.

Figures 8 and 9 show the results of the ablation study, with Fig. 8 showing a representative subject from the test set of the Guy’s dataset, and Fig. 9 showing the box plots for each site, averaged over the three subcortical regions. It can be seen from the example segmentations that the *Standard UNet* failed to segment all of the regions and even the addition of the targeted dropout, whilst it led to a small amount of the third region being segmented, did not lead to a substantial improvement in the performance of the network. Therefore, in this low-data regime and for a relatively difficult task, where the *Standard UNet* failed to complete the segmentation to a standard that would be acceptable, the pruning process alone created a network that is significantly better at segmenting the three regions. Across all folds, the *Standard UNet* found a local minimum which led to one of the regions being segmented very well at the cost of another not being segmented at all. This was also the mode of failure of the pruned network when the amount of training data was reduced to a very low number of subjects (see below). The region that was not segmented does, however, vary between folds and was not necessarily the smallest region.

**Figure 8:**
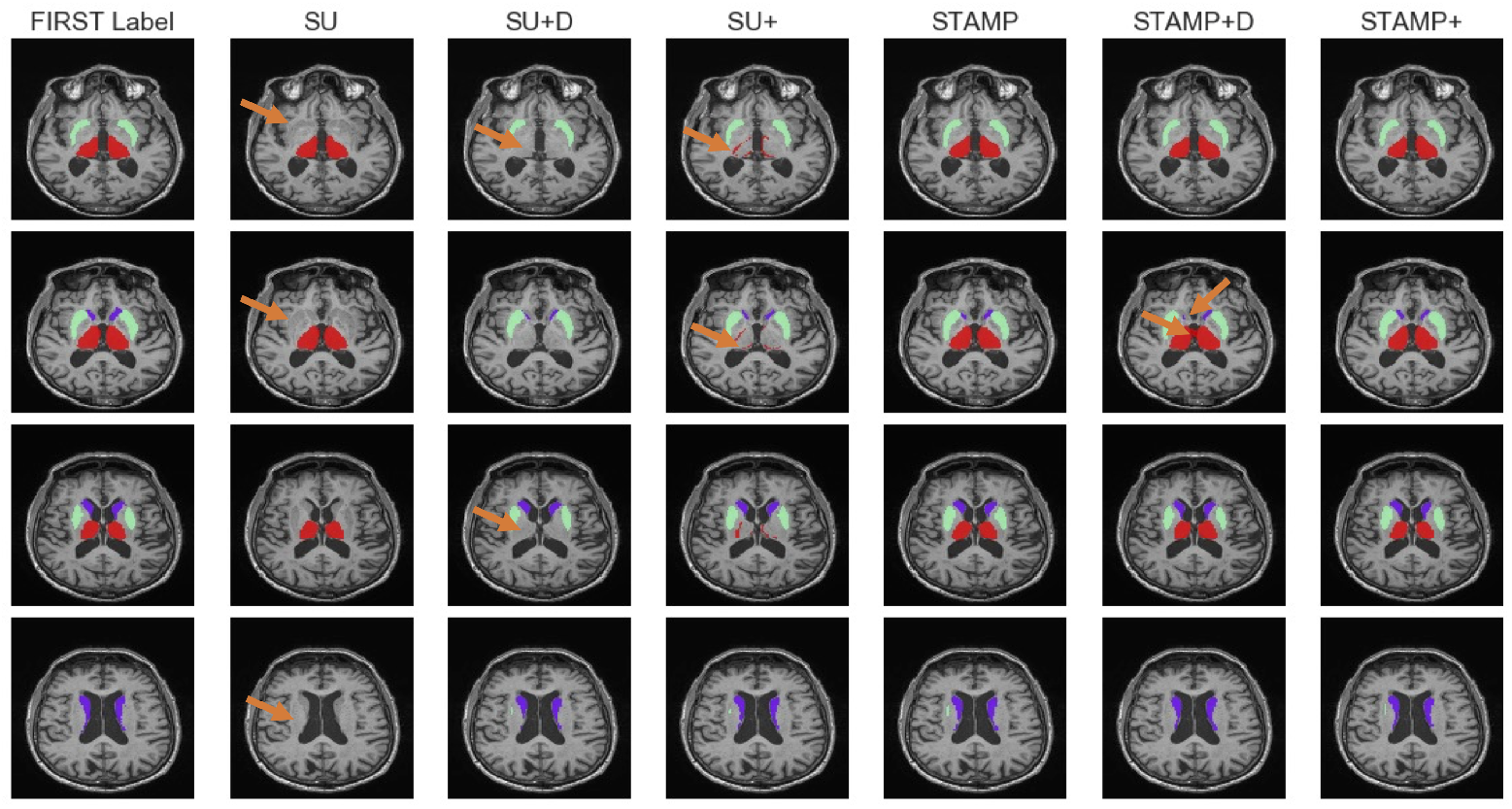
Example segmentations for the caudate, thalamus and putamen trained using the Guy’s data from the IXI dataset for, left to right, starting at the second column: *Standard UNet* (SU), *Standard UNet* with channelwise dropout (SU + D), *Standard UNet* with targeted dropout (SU+), *STAMP*, *STAMP* with channelwise dropout (*STAMP+D*), and *STAMP+*. The labels generated using the FSL FIRST tool (shown in the first column) are used as a proxy for manual labels. Arrows indicate regions with clear differences between approaches.

**Figure 9:**
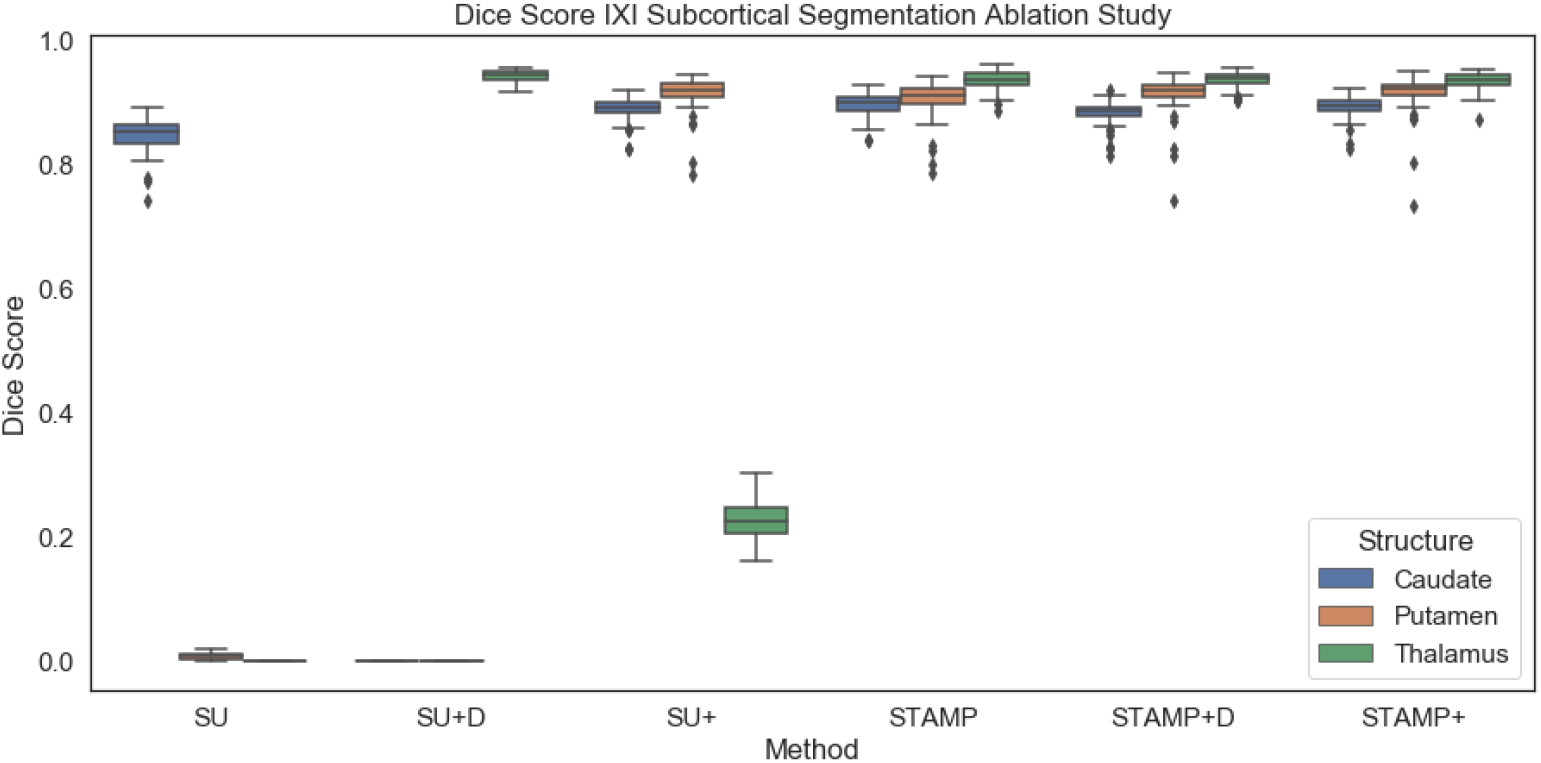
Dice results for the IXI subcortical segmentation ablation study, comparing *Standard UNet* (SU), *Standard UNet* with channelwise dropout (SU+D), *Standard UNet* with targeted dropout (SU+), *STAMP*, *STAMP* with channelwise dropout (*STAMP+D*) and *STAMP+*. It can be seen that even the simplest pruned model, *STAMP*, outperforms all of the models trained without pruning, which were unable to segment all three labels, as seen in Fig. 8. *STAMP+* slightly but significantly outperformed the other approaches for the Dice score averaged over the three regions.

It can also be seen that while the improvement provided by the targeted dropout was relatively small, it was significant (*p* = 0.0007) and, in addition, increased the stability of the *STAMP+* procedure. There was also a significant improvement compared to using a fixed amount of spatial dropout across the networks (*p* < 0.001): this might be expected as this strategy did not encode information about which filters were the most likely to be removed, so there was a chance that filters that were very important to the performance would be dropped, especially as the pruning progressed, hindering the model training. The value of spatial dropout used was very low (0.037), calculated as the average *p* value from *STAMP+*. Network performance was further hindered at higher dropout values.

The poor performance of the *Standard UNet* on this task was almost certainly due to the shortage of training data available for the task. Access to more data for training was not an option in this instance, and so how much more data would be required to improve the performance could not be explored. However, reducing the amount of training data to explore the performance of the *STAMP+* method could be explored. Figure 10 shows the Dice scores on the testing set, averaged across the three subcortical regions. The results broken down by subcortical region for the *Standard UNet* and *STAMP+* can be found in the Supplementary Material. Both sets of results show that the pruned network could be successfully trained with substantially less data than the *Standard UNet* and, in all cases, provided a better performance. This is not just a function of the reduced number of parameters, as we were unable to train a *Standard UNet* with *f* = 2 on the data. Some of the increase in performance might have been due to the network learning coadapted relationships between layers and overfitting less to the very small amount of training data available. Therefore, *STAMP+* clearly provides advantages for segmentation tasks when working in low-data regimes, as is very common in medical imaging applications.

**Figure 10:**
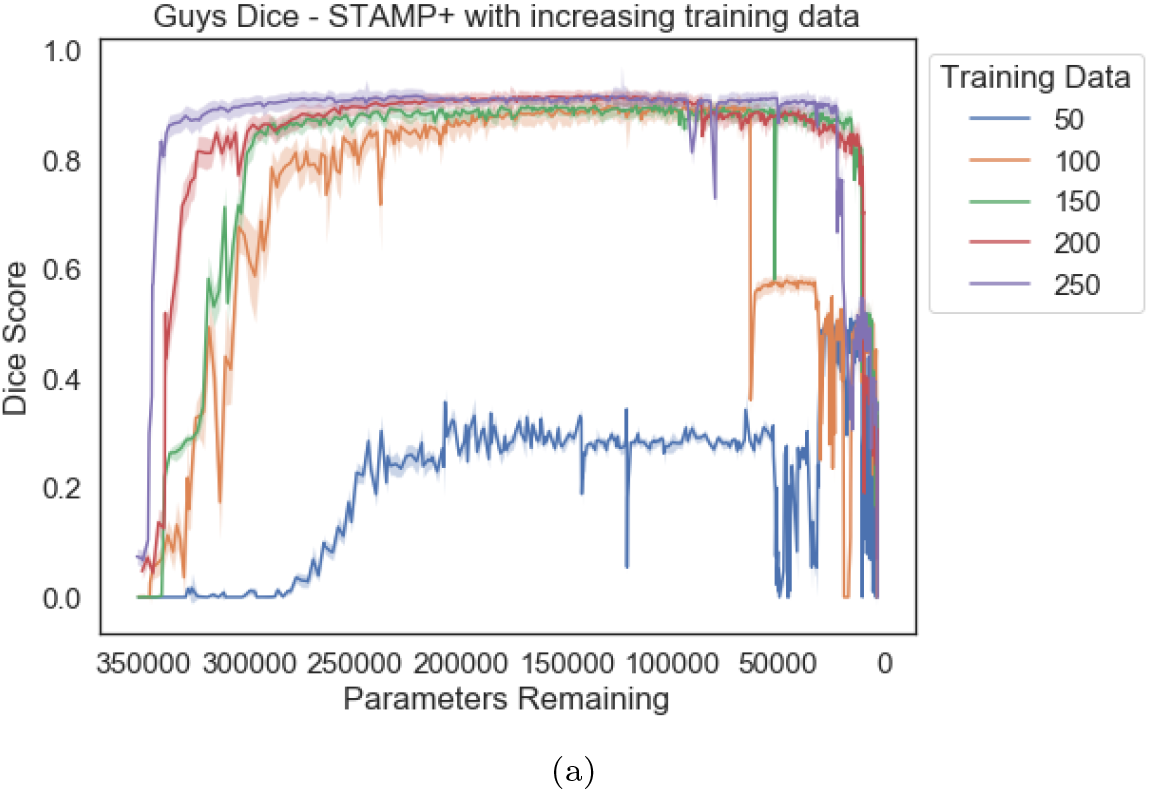
Dice scores averaged across the three subcortical regions for increasing numbers of available training images for the IXI data. 250 represents the full training set. It can clearly be seen that more pruning iterations were required to reach the same dice performance as the amount of data was decreased, until there was insufficient data available.

### 3.3. Comparison of Pruning Strategies

We then compared *STAMP+* to alternative methods for producing smaller models. Namely, comparing it to (i) *PruneFinetune* (section 3.3.1), (ii) changing the initial number of filters in the model architecture *f* (section 3.3.2), and (iii) training the pruned architectures from scratch with normal training (section 3.3.3). We explored these comparisons for the HarP dataset, but the results were seen to be consistent across the datasets considered.

#### 3.3.1. PruneFinetune Comparison

*STAMP+* was therefore first compared to *PruneFinetune* for increasing numbers of training subjects. In Fig. 11 the comparison can be seen for 25, 50 and 100 training subjects. We found that when the number of training subjects was very low (for instance, consider 25 training subjects) the original *Standard UNet* model performed badly on the data. As this model was then the initialisation for the *PruneFinetune* pruning and there were still very few subjects available for training, the model struggled to perform well from the poor starting point. The *STAMP+* model had noisier performance between pruning iterations, but significantly outperformed the *PruneFinetune* approach (*p* < 0.001 for all amounts of training data). Even as the number of training subjects increased, *STAMP+* outperformed *PruneFinetune*.

**Figure 11:**
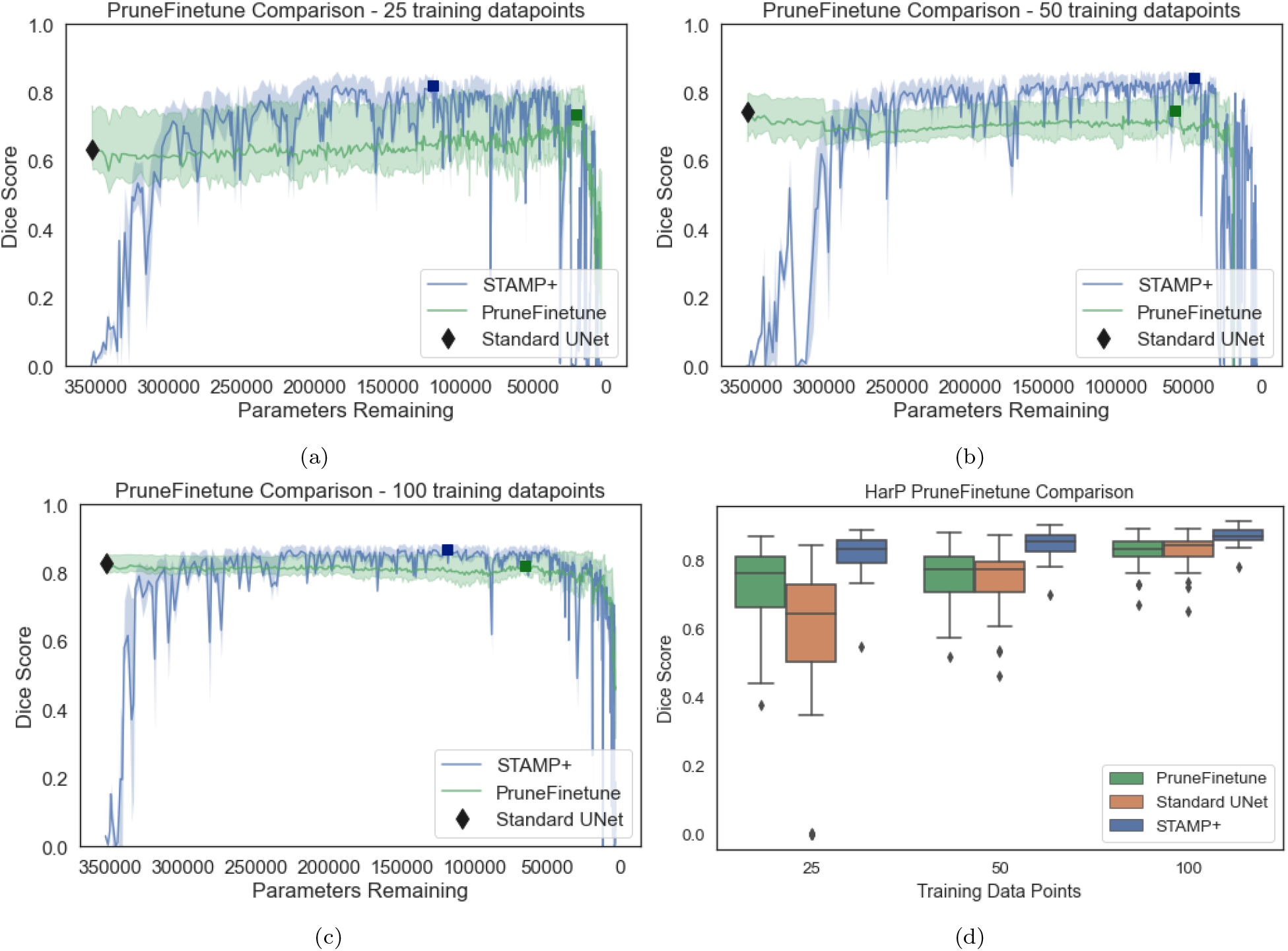
Dice scores for the HarP hippocampal segmentation comparing *STAMP+* and *PruneFinetune* for a) 25, b) 50 and c) 100 training subjects. Although *PruneFinetune* is more stable between iterations than *STAMP+* for low data regimes, its performance is significantly worse and produces inconsistent predictions. Even as the amount of data is increased, *STAMP+* significantly outperforms *PruneFinetune*. d) The boxplots show the best performing model (selected on validation data) for *PruneFinetune* and *STAMP+* compared to the *Standard UNet*. The models selected are indicated on subplots (a-c) by the correspondingly coloured squares.

This result was for *PruneFinetune* with only a single filter being removed at a time, which can be regarded as very conservative compared to the normal approaches in the literature, where a percentage of filters, or all filters under a threshold value are removed (Cun et al., 1990), which we would expect to be less stable. Furthermore, the recovery time between filter prunings was much longer than that of *STAMP+*, as the model was allowed to recover entirely before additional pruning is performed (representative training graphs can be found in the Supplementary Material). Finally, the original model had to be trained to convergence before the network could be pruned, and so *PruneFinetune* had a much larger computational cost while performing less well than the proposed method of *STAMP+*. Therefore, the proposed method allowed better performance to be achieved without having to train the original model.

Ideally, these results would be repeated across all the datasets, however, the amount of computational time required to run these experiments would be vast. The number of epochs could be reduced by increasing the pruning percentage per pruning episode, but we found in preliminary results that the lower the percentage, the better the performance, as the network is able to recover from the pruning episode best. Also, we found that pruning a single filter at a time gave the best Dice results, and so is a fairer comparison to our proposed *STAMP* method. Thus, for each of the remaining datasets, we compared *STAMP+* to *PruneFinetune* for the lowest amount of training data considered, as this low data regime was where *STAMP+* showed the most improvement over the *Standard UNet*. The results can be seen in Fig. 12, where it is evident that *STAMP+* consistently outperforms *PruneFinetune*, most likely due to the poor initialisation of the networks from the *Standard UNet* models, due to the limited training data. For all but the prostate dataset, *PruneFinetune* outperformed the *Standard UNet*, showing that pruning was generally useful for low data model training, but the additional improvement from *STAMP+* showed the benefit of training and pruning simultaneously.

**Figure 12:**
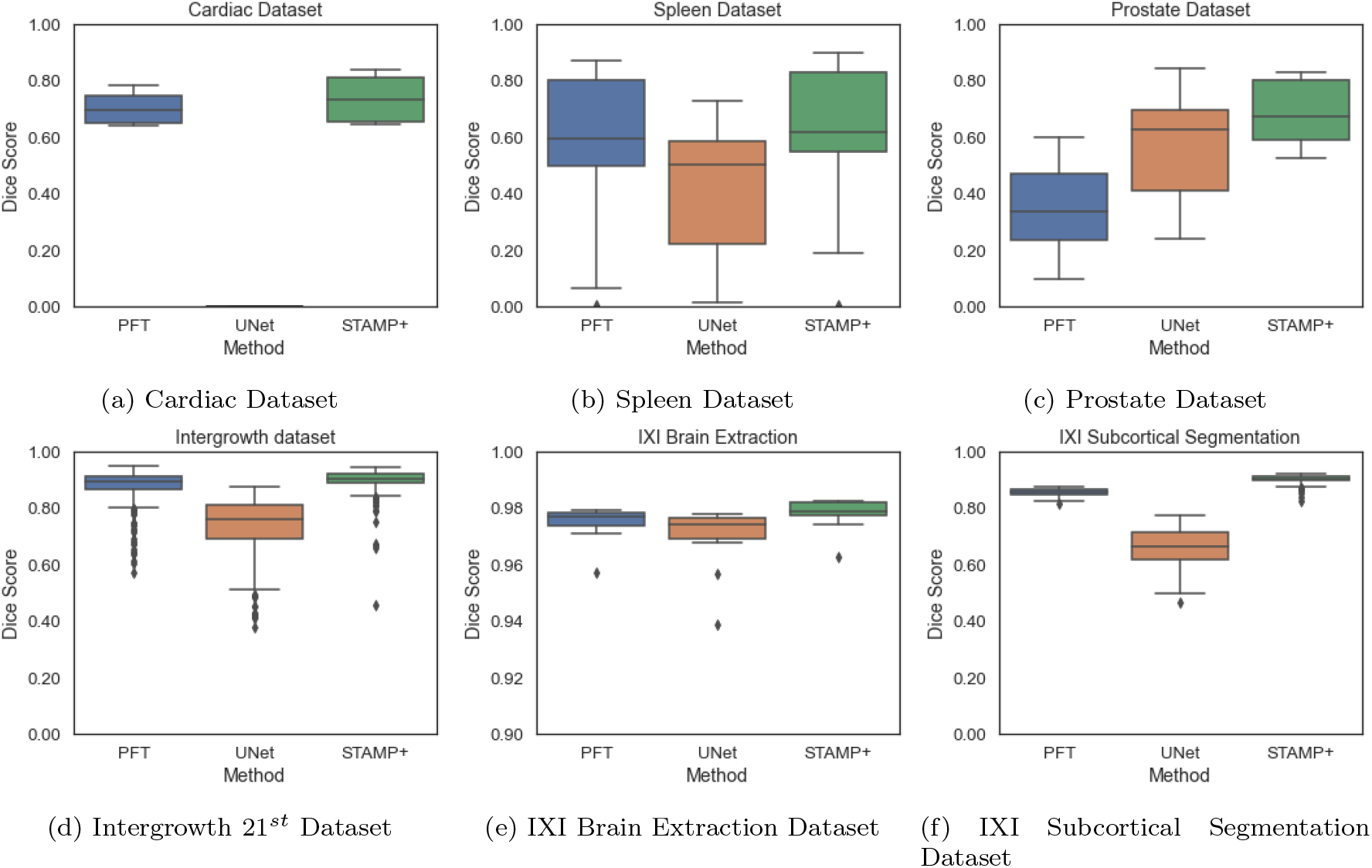
Comparison of the *Prunefinetune* (PFT), *Standard UNet* (UNet) and *STAMP+* across the datasets explored, for the smallest number of data points explored for each dataset. Note the different y-axis scales, to enable the methods to be compared.

#### 3.3.2. Initial Number of Filters

We then considered changing the initial number of filters, *f*, present at each depth of the model. Changing the initial number of filters represents a different way in which the size of the model can be modified other than through model pruning. Figure 13 explores training with increasing numbers of training data points for models with *f* = {2, 4, 8} compared to the *Standard UNet*. It is evident that for all sizes of model and for all amounts of available training data, *STAMP+* outperformed the *Standard UNet*. Further, the performance of the *Standard UNet* was worst when using *f* = 2, indicating that naïvely reducing the number of parameters was detrimental to model performance. Through pruning, *STAMP+* was able to create models of the same size or smaller than the naïve *f* = 2 UNet, while outperforming them in terms of Dice score. Further analysis of this performance can be found in the Supplementary Material.

**Figure 13:**
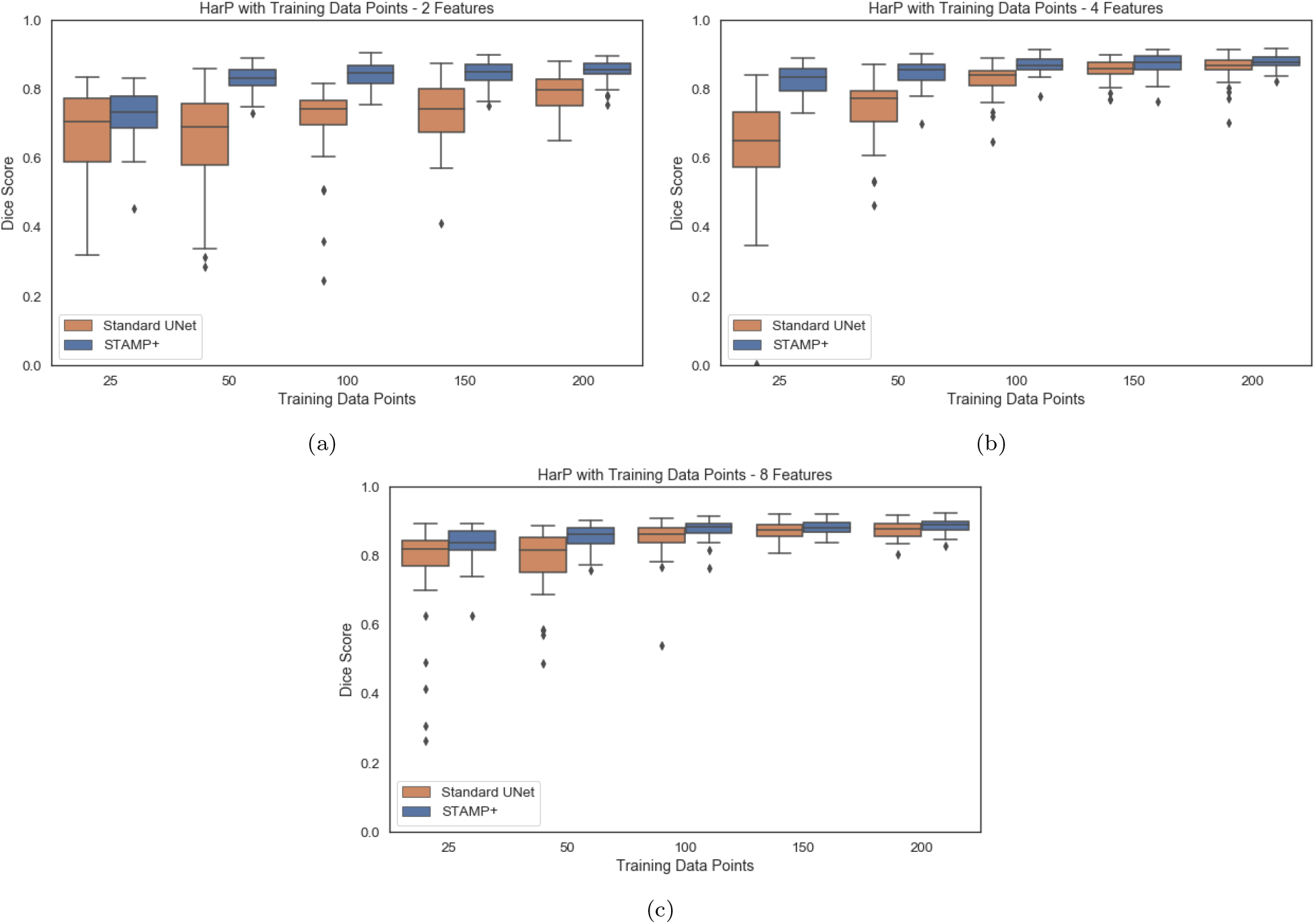
Dice scores for the HarP hippocampal segmentation for increasing amounts of training subjects, comparing *STAMP+* and the *Standard UNet*, for *f* = {2, 4, 8}.

Figure 14 shows the distribution of filters of the differently sized networks both at initialisation and at the point in pruning where they had the same number of parameters as the *f* = 2 model, when training with the full HarP dataset. The number of filters in these pruned models corresponded to the *f* = 2 model trained to convergence, which achieved a Dice score of 0.788 ± 0.052 compared to the pruned models’ performances of 0.833 ± 0.041 and 0.859 ± 0.047 for *f* = 4 and 8, respectively. This means that all of the larger pruned models performed significantly better than the *Standard UNet* model of that size. It is clear that the filter distributions learned through pruning the models were very different to the *Standard UNet* architecture of doubling filters, instead having far flatter distributions, with a similar number of features at each depth being maintained. A shallower network architecture initially was also considered, which led to the same pattern of results, and can be found in the Supplementary Material.

**Figure 14:**
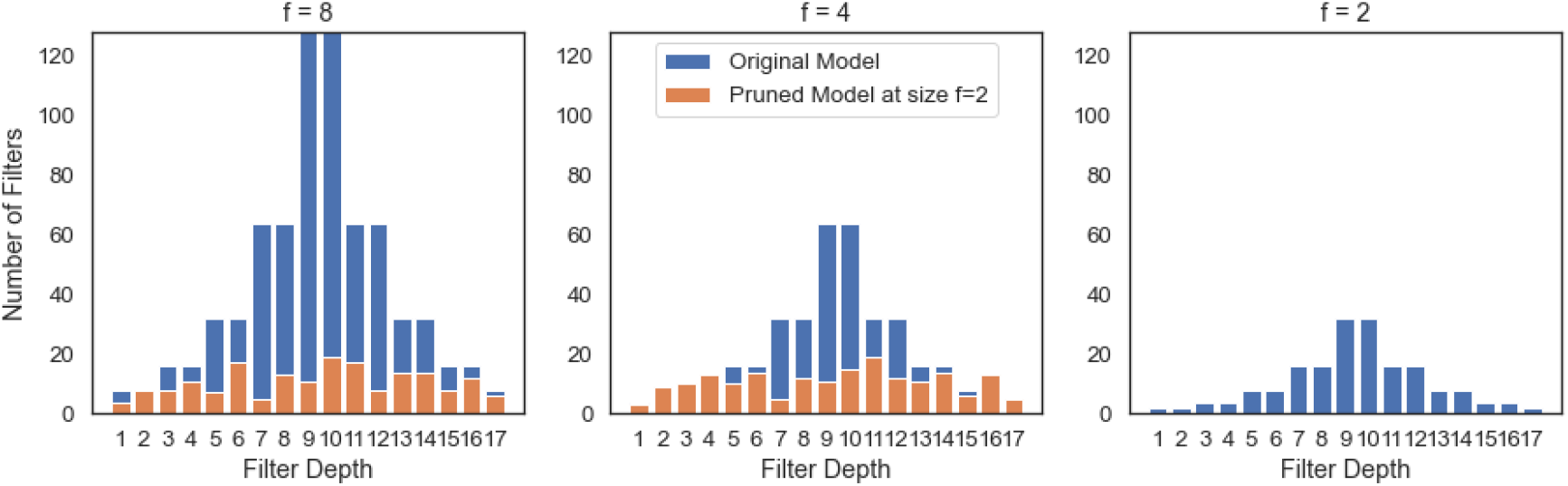
The number of filters in each layer for the models trained on the full HarP dataset in Fig 13. The distribution of the number of filters shown in blue corresponds to the model’s original filter distribution, and the one shown in orange is the distribution when the model has the same total number of parameters as the original model with f = 2.

#### 3.3.3. Training Pruned Models from Scratch

Having demonstrated in the previous experiments that naïvely reducing the size of the network is not sufficient to improve model performance and to ensure that the model improvement seen is a function of the simultaneous training and pruning that characterises our method, we take the intermediate network architectures and retrain them from random initialisation. Figure 15 compares the performance of a single run of *STAMP+* with a set of *Standard UNets* of decreasing network size, using the full HarP dataset for training. For *STAMP+* the results for every 40^*th*^ pruning iteration are shown, and each one is used to define a network architecture that is then trained as a *Standard UNet*. For the first model size – full model architecture – the performance of *STAMP+* is poor as the network has only had a single epoch of training. It can be seen that, once *STAMP+* has completed sufficient epochs of training, STAMP+ outperforms the equivalent UNet model (in terms of architecture) when they are trained from scratch, especially as the networks become very small in terms of numbers of parameters. Therefore, the model performance is a function of both the smaller architecture and the simultaneous training and pruning, possibly due to the fact that *STAMP+* learns co-adapted features which cannot be directly learned when training the smaller network architectures.

**Figure 15:**
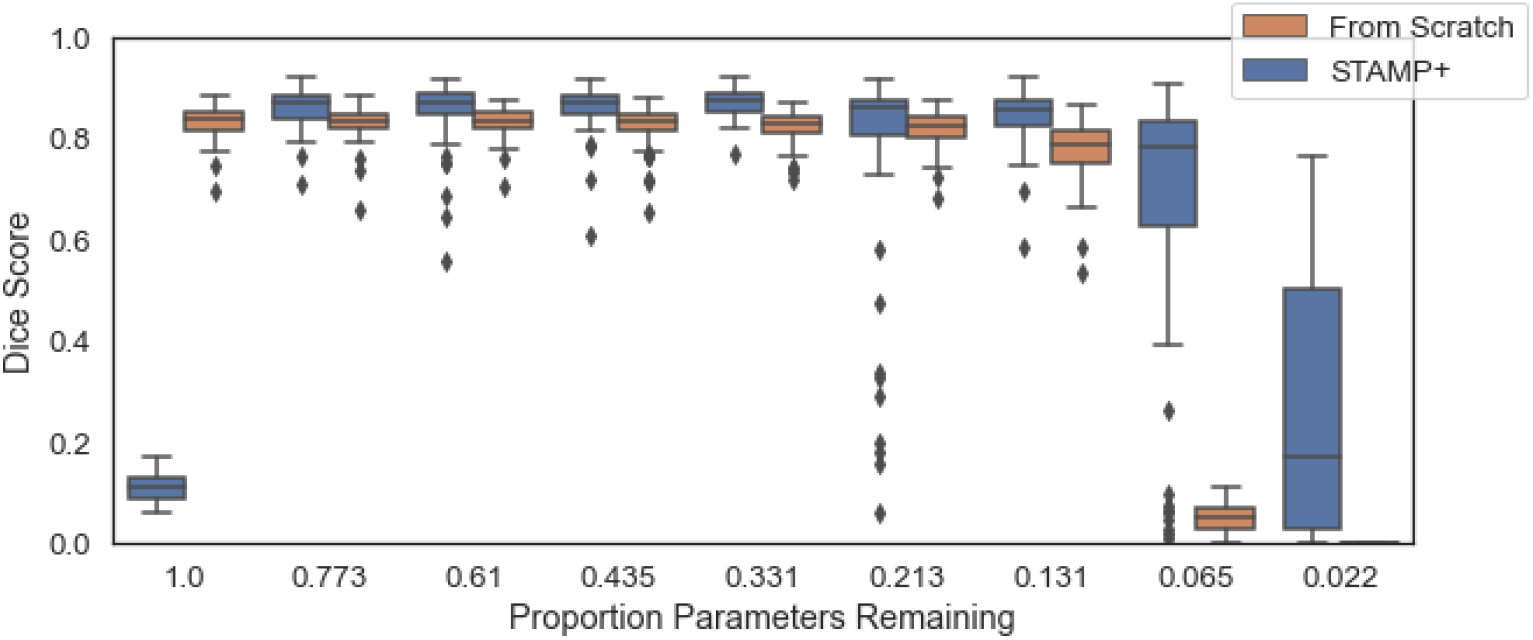
Comparison of the results from *STAMP+* to training the model architectures identified by *STAMP+* from scratch on the HarP data, having been randomly reinitialised. The results are shown for every 40^*th*^ parameter, with the number of parameters remaining in the model shown. Note that for the largest size model, *STAMP+* had completed a single epoch of training, and so the performance is very low.

### 3.4. Effect of Hyperparameters

*STAMP+* introduced two new hyperparameters into the training process, the effect of which need to be understood: the number of recovery epochs between pruning episodes, and the *b_drop_* value which controlled the magnitude of the targeted dropout values. These will again be explored using the HarP dataset.

#### 3.4.1. Recovery Epochs

*STAMP+* requires iterating between training the network and pruning filters one at a time. We therefore explored the effect of the number of recovery epochs between filter prunings, when the network was allowed to train without pruning. Figure 16 shows the performance of the network with pruning for increasing numbers of recovery epochs, when 50 training data points were available. The results for other amounts of training data can be found in the Supplementary Material, where we have shown that the pattern was the same across the amounts of data, but the impact of increasing the number of recovery epochs decreased as the amount of data available for training increased.

**Figure 16:**
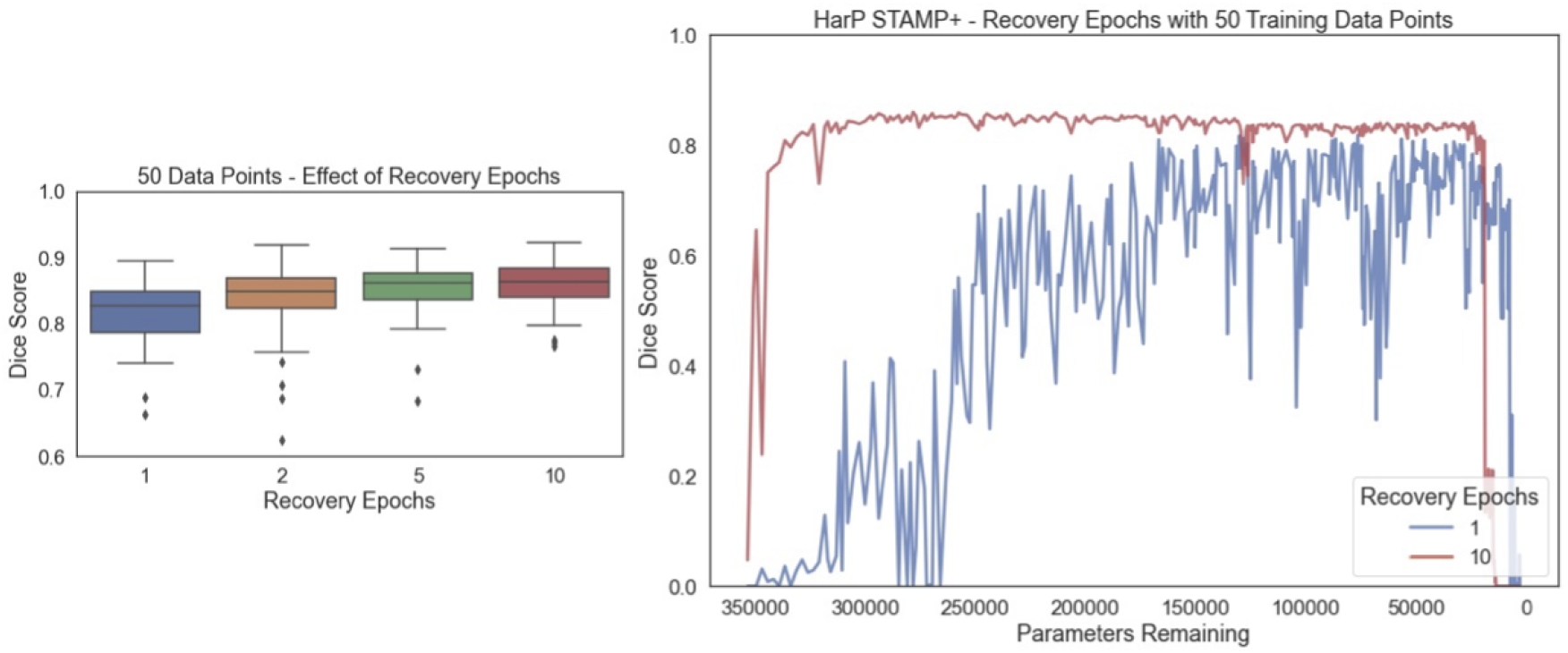
Effect of increasing the number of recovery epochs on the segmentation performance on the HarP data with 50 training examples. The box plot shows the best performance achieved, chosen based on the validation data, and the lineplot shows the pruning Dice score for 1 and 10 recovery epochs as the training progresses, showing the mean value as the number of parameters decreased, clearly showing the increase in stability and performance for increasing the number of recovery epochs. The pattern was the same across the other amounts of training data, as can be seen in the supplementary material.

It can be seen that the maximum performance reached was similar in all cases (Dice: 1*ep* = 0.817, 2*ep* = 0.836, 5*ep* = 0.853, 10*ep* = 0.860, with the best performing model selected using the validation data), but improved with increasing recovery epochs. The degree of improvement was more evident with lower amounts of training data. The biggest difference between runs as the recovery epochs were changed was the stability of the training, with the stability in performance between pruning iterations increasing with the number of recovery epochs. Therefore, the setting of the recovery epochs was a trade-off between the stability and the duration of the training.

#### 3.4.2. Base Dropout Value *b_drop_*

Finally, we considered the effect of the *b_drop_* value, which modulated the values of the targeted dropout. Figure 17 explores the effect of changing the value of *b_drop_* on the proposed *STAMP+* method. Across the experiments, a base value of 0.10 was used, meaning that the maximum dropout probability that could be applied to a layer is 0.10. This is a lot lower than the dropout probability used with parameterwise dropout (Srivastava et al., 2014) where values as high as 0.5 are regularly used; however, spatial or channelwise dropout is much more aggressive. This value however led to a significant increase in performance as can be seen by comparing *STAMP* to the addition of targeted dropout with *b_drop_* = 0.10 (*p* < 0.001) in Fig 17. Very similar performance was observed for a large range of dropout values 0.05-0.30, showing that the method was not too dependent on the value chosen. As the *b_drop_* value is increased to very high values, 0.40-0.50, it can be seen, as would be expected, the performance degraded and the training became very unstable, due to the large amount of parameters being dropped out.

**Figure 17:**
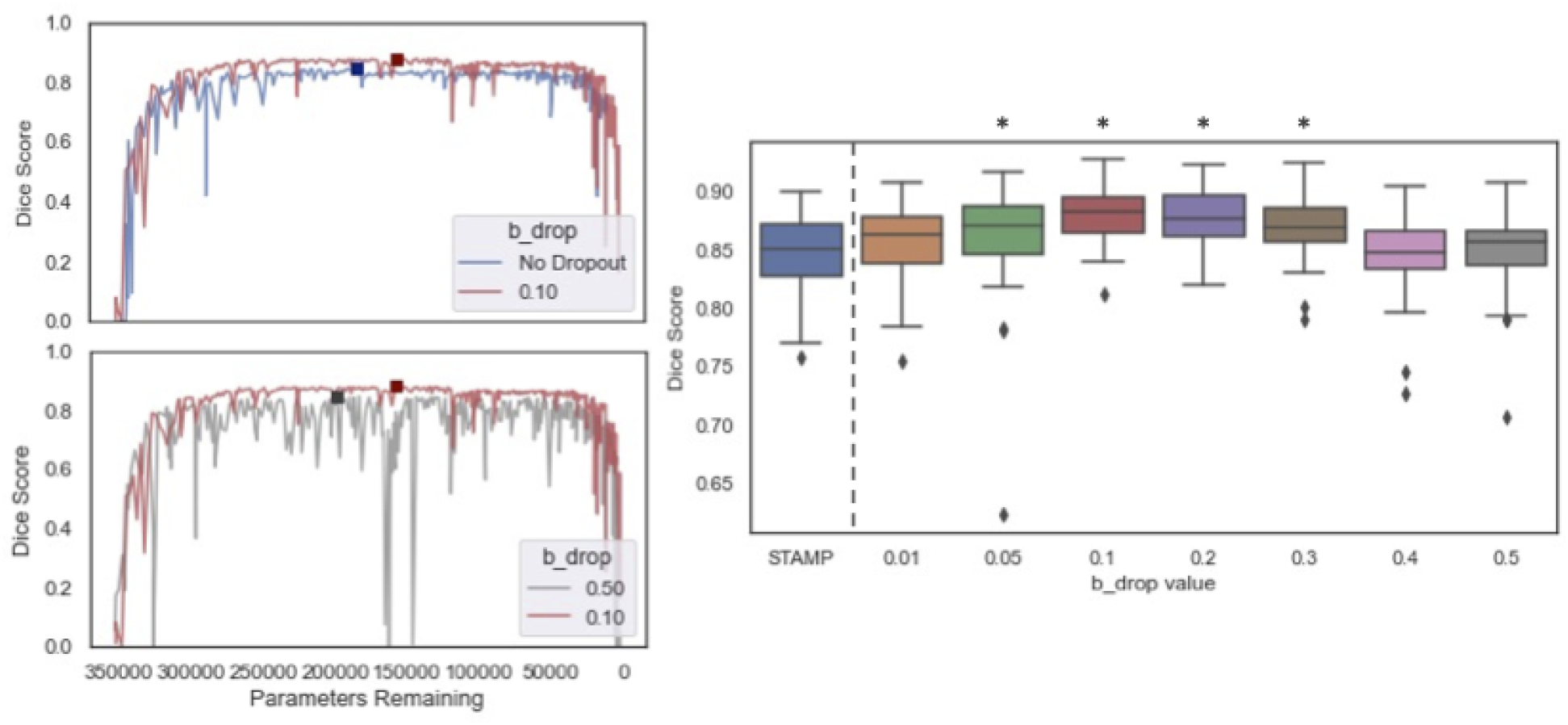
The effect of *b_drop_* on the model pruning. The line graphs show the Dice value against the number of parameters remaining comparing no dropout (*STAMP*) to *STAMP+* with the value of *b_drop_* used (0.1), and this *STAMP+* configuration (*b_drop_*=0.1) to *STAMP+* with a much higher *b_drop_* value. The change in stability of *STAMP+* from changing the *b_drop_* value is clearly visible, and an increase in performance can be seen compared to STAMP with no dropout. The boxplots then show the maximum dice scores achieved for the HarP testing data, with increasing *b_drop_* value. * indicates significant improvement over *STAMP* baselines, *p* < 0.001. Note: no dropout corresponds to *STAMP*.

Therefore, the results were sensitive to the choice of the *b_drop_* value, but there was a large range of values for which good performance was achieved and there was clear benefit from its addition, highlighting the merits of *STAMP+* over *STAMP* alone.

## 4. Discussion

We have presented a method that performs simultaneous training and pruning of a UNet architecture for medical image segmentation. While pruning has been well explored for large classification datasets, to the best of our knowledge it has not been explored for medical image segmentation, where different architectures and generally small training datasets create a different challenge for pruning algorithms. Therefore, the *STAMP+* method and the addition of Targeted Channelwise Dropout have been explored for a range of segmentation datasets and tasks: we have shown that not only is it possible to create smaller models with as good or increased performance on the task of interest, but also that, through pruning, better performing models can be produced in low data regimes.

Using the HarP data, the method has been validated on data with manual segmentation labels and the effects of the design decisions have been explored. The results indicate that better performance is achieved by initially choosing a large model and pruning the model to be smaller, rather than beginning with a small model, and that different distributions of the filters are kept by the pruning process than might be naïvely chosen. The experiments also show that the addition of the Targeted Dropout led to an improvement in performance and that this improvement was present for a relatively wide range of *b_drop_* values (0.05 to 0.30).

It has been demonstrated that the hyperparameters used are robust across a range of tasks and imaging modalities, for both 3D and 2D problems, showing that the method is applicable across a range of medical imaging tasks, and may be used in many cases without the need for extensive parameter searching or optimisation. That is, the default parameters led to the same or improved performance compared to the *Standard UNet* for all cases. Furthermore, the increase in performance in low data regimes has been demonstrated across these different datasets and tasks. The ablation study on the IXI dataset indicated that the improvement is not just a function of the pruning or the targeted dropout, but a combination of the two aspects.

An alternative for low data segmentation that could be considered is the use of atlas-based methods. Atlas-based methods have been shown to produce good quality segmentations in low data regimes (Lee et al., 2019); however they are limited to cases where atlases are available. If a small number of labels were available, they could be used to construct an atlas. However, recent results suggest it is better to use the labels alongside augmentation to train CNNs directly (Hesse et al., 2022). Further, there are segmentation tasks where atlases cannot be constructed meaningfully, for instance white matter hyperintensities, which are not suited to atlases as they vary so much in terms of location and size. Thus, our proposed method offers a more general solution to segmentation in very low data low label regimes.

Whilst care has been taken to test the method across modalities and segmentation tasks, the labels used for the IXI dataset were generated using automated tools. The two tools used – FSL FIRST and ANAT – employ differing model assumptions so the results should be valid for comparison between methods; however, it does potentially limit the maximum achievable performance due to the imperfections in the labels.

The method has also only been explored for the standard UNet architecture. This decision was made as the UNet is the most popular architecture for medical image segmentation, and the majority of methods either use the UNet architecture or derivatives thereof. It is expected that the results would generalise to other similar networks, but this has not been explored explicitly within this work, and so future work should focus on exploring the approach for other network architectures commonly used in medical imaging.

Finally, a potential limitation of the method is that the performance between pruning iterations is unstable, especially compared to the *PruneFinetune* method. Across this work, the best model was evaluated using the validation data, and this has corresponded well with good performance on the testing data – although it does not always correspond to the highest performing iteration on the testing data. This has been true across the datasets explored here but would not necessarily be true if the method were to be applied to other datasets. The use of the targeted dropout helps to reduce this by increasing the stability between iterations, but if this were not seen to be the case for a given dataset, it may be necessary to increase the number of recovery epochs to increase the stability of the model training. It is probable that a lot of the instability is due to working in the low data regime, as the stability visibly increases as the number of training subjects is increased. In practice, a large number of recovery epochs could be used, and gradually reduced to smaller numbers if the training were stable, and could be simply automated.

## 5. Conclusion

We have developed and demonstrated a method to allow simultaneous training and pruning of a UNet architecture. We have shown that in low data regimes this outperforms the equivalent *Standard UNet* models and the standard pruning method due to removing the need to train the original full-sized model. The channelwise targeted dropout assists the pruning, by making the model more robust to being pruned. The method has been demonstrated across different organs, segmentation tasks and imaging modalities and it is expected that the results should generalise to other network architectures.

## 6. Data and Code Availability Statement

The data used in these experiments are available from the relevant studies. The code is available at github.com/nkdinsdale/STAMP. Weights from training are available on request through emailing the corresponding author.

## 7. Acknowledgements

ND is supported by the Engineering and Physical Sciences Research Council (EPSRC) and Medical Research Council (MRC) [grant number EP/L016052/1]. MJ is supported by the National Institute for Health Research (NIHR), Oxford Biomedical Research Centre (BRC), and this research was funded by the Wellcome Trust [215573/Z/19/Z]. The Wellcome Centre for Integrative Neuroimaging is supported by core funding from the Wellcome Trust [203139/Z/16/Z]. AN is grateful for support from the UK Royal Academy of Engineering under the Engineering for Development Research Fellowships scheme.

The computational aspects of this research were supported by the Wellcome Trust Core Award [Grant Number 203141/Z/16/Z] and the NIHR Oxford BRC. The views expressed are those of the author(s) and not necessarily those of the NHS, the NIHR or the Department of Health.

## 9. Supplementary Material

### 9.1. Choice of Metric

Here, we explored the choice of metric used to determine which filters to prune. As stated in section 2, the L2 norm was chosen as the pruning metric and the results shown in Fig. 18 formed the basis of that decision. For N data points, the L2 norm was compared with:

- L1 norm (Li et al., 2017): 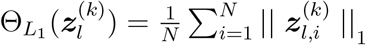
- Taylor – proposed in (Molchanov et al., 2016), which utilises the product of the activation and the gradient of the loss function: 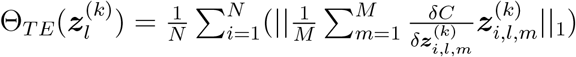 where *m* is the element in the vectorised feature map of length *M*.
- Random: Θ_*Rand*_ ~ *Uniform*(0, 1)

where all the metrics but Random are normalised according to Equation 3. This normalisation ensured that the last filter at any depth was not pruned. With Random pruning, the condition that the last kernel at a given depth cannot be pruned was explicitly coded. Figure 18 shows the Dice scores on the test dataset at each pruning iteration for models trained using *STAMP+* and with each metric in turn. For each metric, the mean value across the test set is shown as the solid line, and the shaded region indicates the interquartile range. The results are plotted against the remaining parameters rather than the pruning iteration. This was done because different pruning iterations remove different numbers of parameters, depending on network location from which the filter was removed: for instance, if it also led to filters being removed across the skip connection.

**Figure 18:**
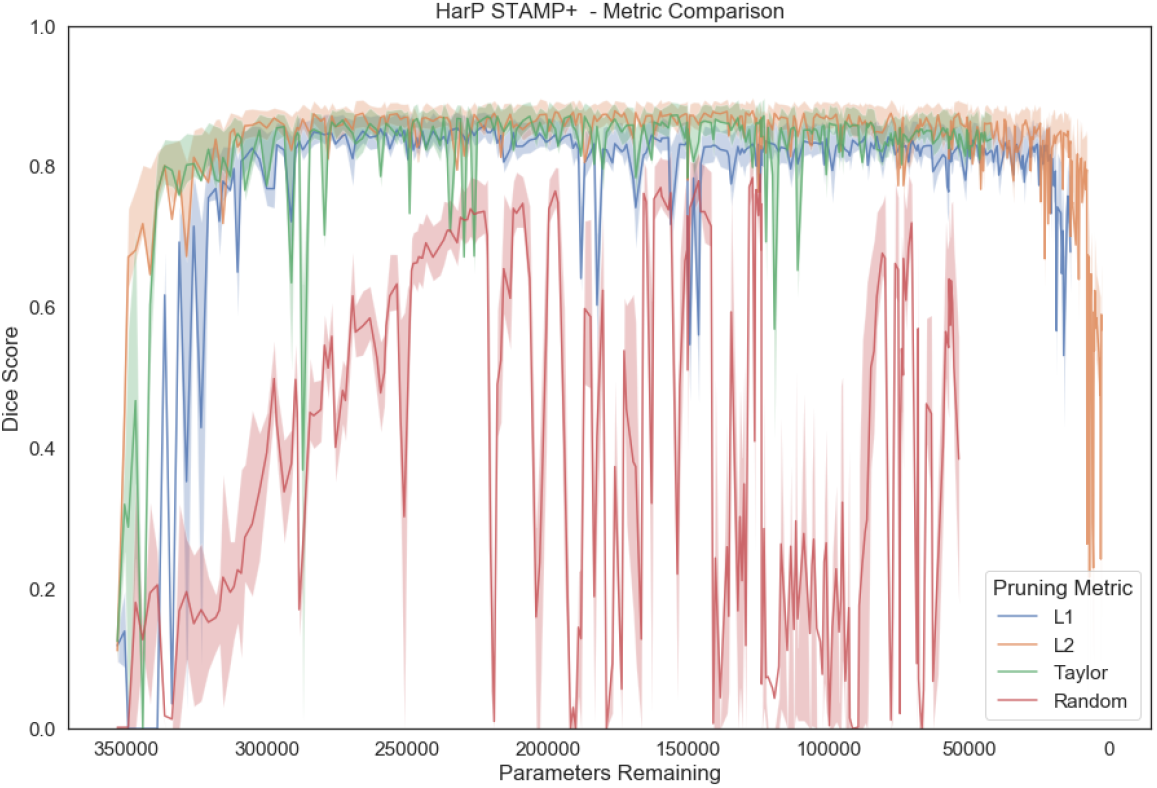
The performance of the *STAMP+* algorithm with different choices of pruning metric using the HarP data, with *f* = 4. It can be seen that the three metrics performed similarly, with no significant difference between L2 and Taylor metrics. All three metrics were better than random pruning. Note that the results are plotted against the remaining parameters rather than the pruning iteration.

It can first be seen that all three metrics performed better than randomly pruning channels, as would clearly be expected. As the models started from random initialisation, the performance initially for all four metrics was poor, then improved rapidly for all metrics except random as the model training continued. It can then be seen that all three of the metrics performed comparably on this task, with no significant difference between the performance of the L2 and Taylor metrics (L2 vs Taylor: *p* = 0.07, L2 vs L1: *p* = 0.003, L2 vs Random: *p* = 5.7 × 10^-11^). As the L2 norm is computationally more efficient than the Taylor metric, it was used throughout the experiments.

### 9.2. Number of filters - Further exploration

The *STAMP+* algorithm was tested on the HarP data for models with varying numbers of filters: *f* = [2, 4, 8, 16]. Each model was simultaneously pruned and trained until the model was unable to be pruned any further, leaving only one filter remaining in each layer. Figure 19a shows the performance of the model: the performance was evaluated on the test set for each iteration of the pruned model; the mean value (the solid line) and the interquartile range bounds (shaded) are shown. The results are shown for four different initial sizes of model, plotted against the number of parameters remaining in the model. For comparison, the correspondingly sized *Standard UNet* models were also trained until convergence.

**Figure 19:**
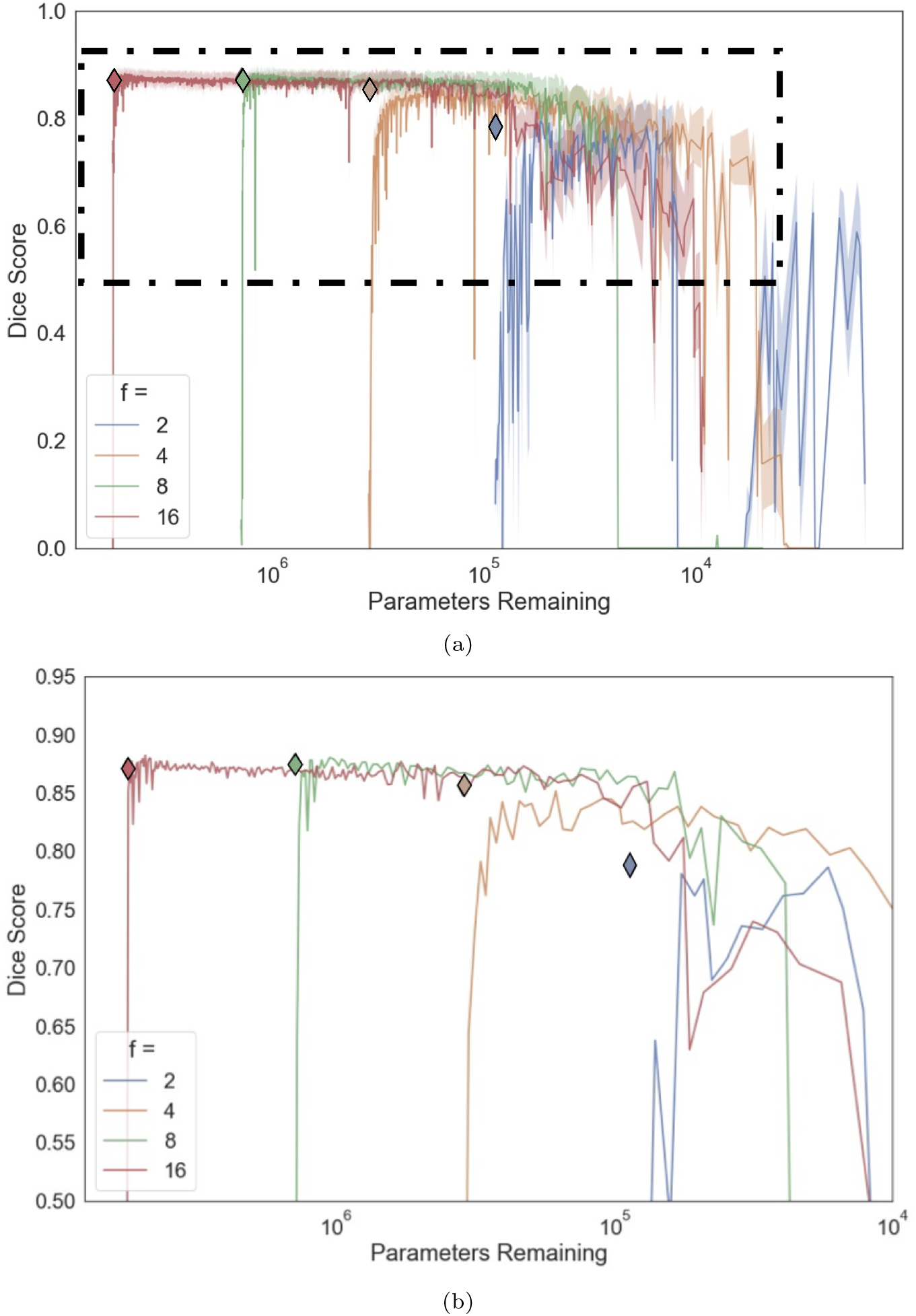
a) Results for pruning models of varying model sizes – using different values of *f* – while training to segment the hippocampus on the HarP data. The black diamond corresponds to a model of the same size being trained to convergence on the same data, with all hyper-parameters held the same except for the absence of pruning the network. b) The same result, zoomed in and subsampled (only every 10^*th*^ data point shown, only the mean, with no interquartile range shown) to allow the result to be seen more clearly. The box on a) indicates the zoomed in region.

First, it can be seen that the model could be pruned and trained simultaneously, such that the same network performance was reached as the model trained to convergence; therefore, the pruned network was sufficiently powerful to be able to represent the variation in the data. It is also evident from Fig. 19 that it was possible to prune the models to a fraction of their original size without reducing the network performance. We also found that the smallest model, when *f* = 2, was not able to perform the segmentation well (maximum Dice score = 0.788 ± 0.052 for the standard UNet and 0.799 ± 0.050 for the pruned model); however the larger networks could be pruned to the same number of parameters as the *f* = 2 model (and smaller) and still perform well on the segmentation task. This indicates that by pruning a larger model we were able to learn a better arrangement of parameters than when naïvely creating a model of that size.

### 9.3. Shallower Network

To allow further exploration of the effect of the initial size of the model, pruning a shallower original model was also considered. A downsampling and an upsampling layer were removed, so the original model was smaller, with fewer parameters, but still having the ability to learn the same number of features at the layers of abstraction that remained. A shallower network was considered, rather than a deeper one, as it allowed the exploration of another way in which the model could be smaller in terms of parameters. This architecture can be seen in Fig. 20. All other parameters were held the same as in the experiments with the full-sized model and the value of *f* was varied.

**Figure 20:**
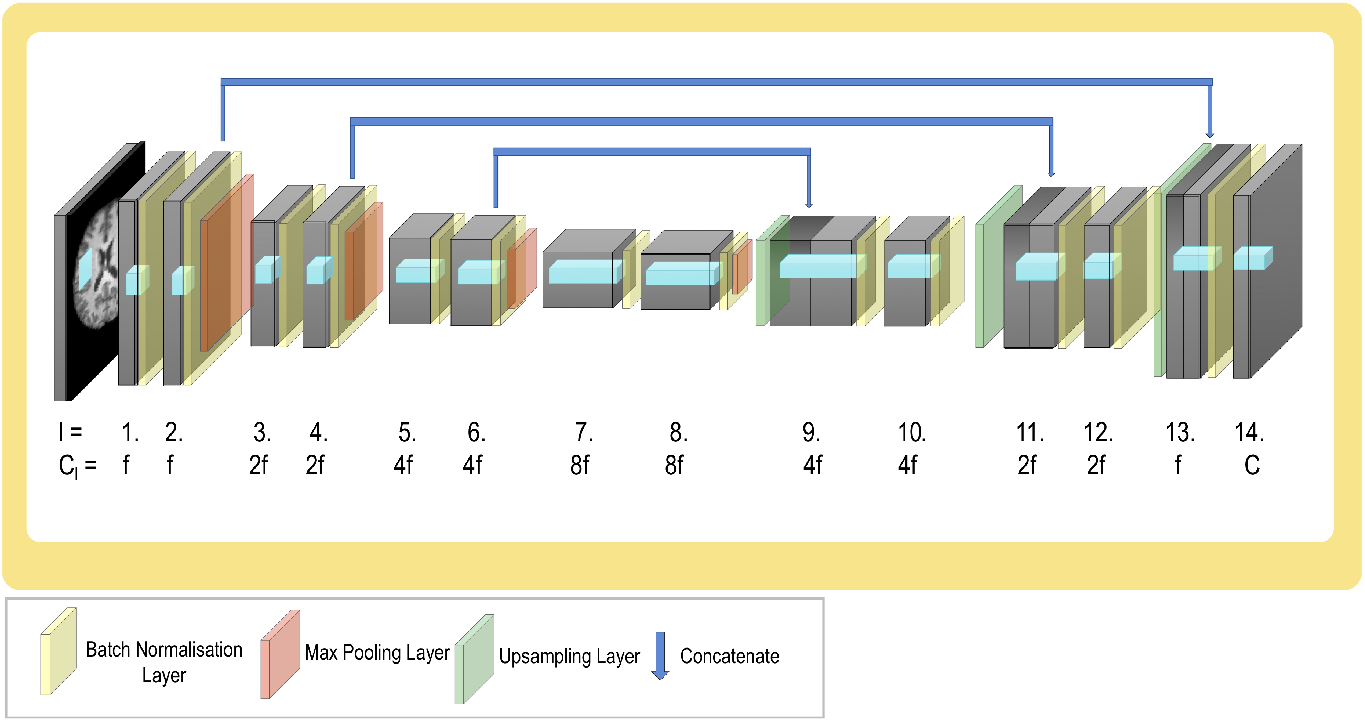
Shallower UNet model architecture - it follows the standard pattern of halving resolution and doubling filters at each depth but has less depth than the UNet architecture previously considered. *l* corresponds to the layer depth, *C_l_* is the number of channels in that layer and *C* is the number of classes in the output segmentation.

We first investigated results obtained using the HarP data, for different values of *f* = [2, 4, 8, 16]. Figure 21 shows the results, where it can be seen that the results with the shallower UNet followed the same pattern as was seen with the *Standard UNet* - Fig 19. Again, it is clearly visible that a better performance on the segmentation task was achieved if we began with a larger model and pruned it to be the same size as the smaller model, rather than originally training the smaller model to convergence. The pruning was, however, less stable, showing that the model training was less robust to filters being removed than when the model was deeper.

**Figure 21:**
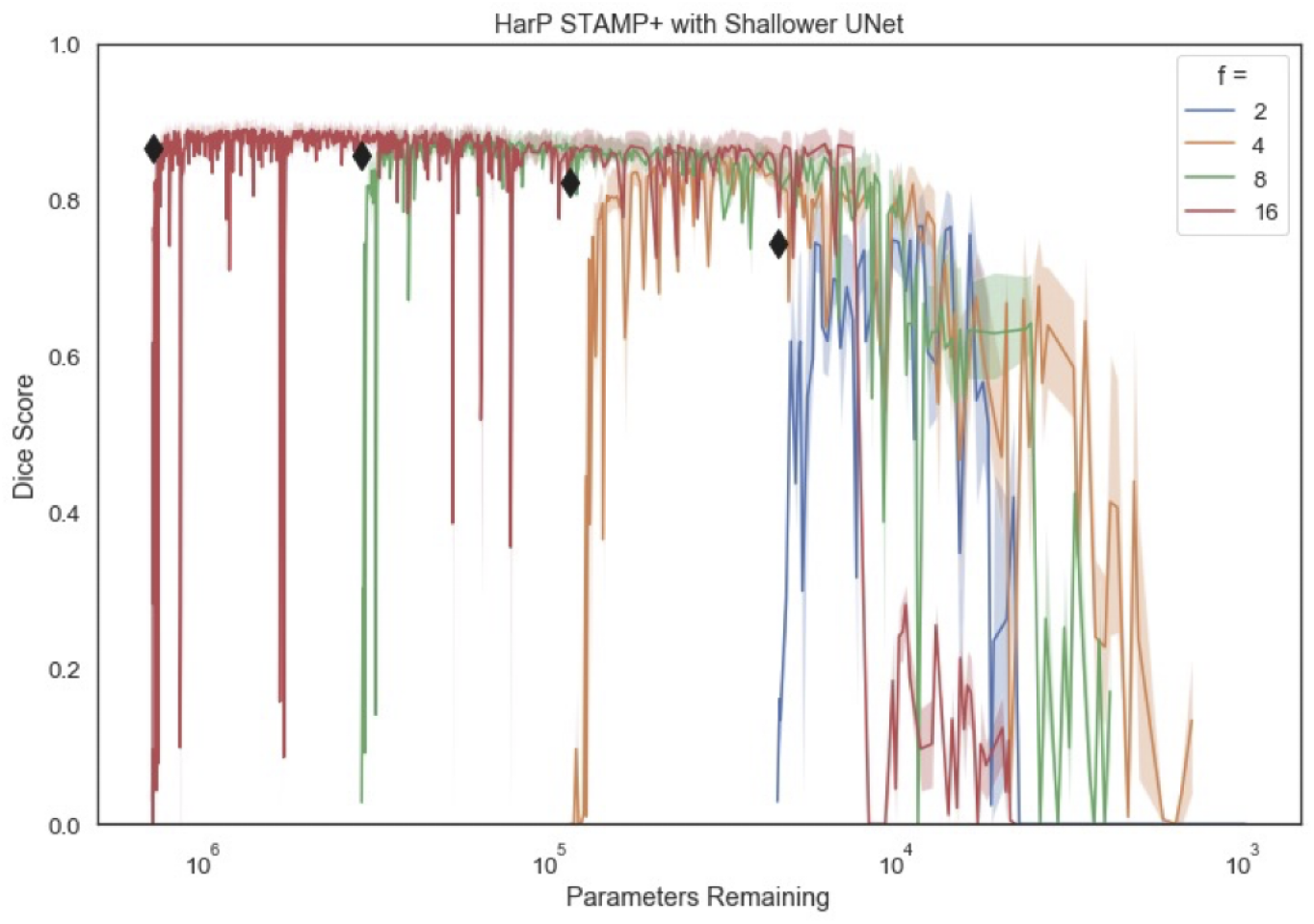
Dice scores on the HarP data using the shallower UNet architecture for *f* = [2, 4, 8, 16], with interquartile bounds shown, using *STAMP+*. Diamonds represent a *Standard UNet* with that number of parameters trained to convergence.

Fig. 22 compares the results from the shallower UNet to the original UNet when the original models were matched in terms of parameters, where Fig. 22a compares an original depth UNet with *f* = 4 to a shallower UNet with *f* = 8, and Fig. 22b compares an original depth UNet with *f* = 8 to a shallower UNet with *f* = 16. The black diamonds represent the original *Standard UNets* trained to convergence; they performed very similarly on the data. As they are pruned, the two similarly sized networks had very similar performance on the main task, indicating that the exact arrangement of the filters is not vital for the model’s performance. This therefore suggests that the algorithm was reasonably robust to the initial model configuration and the results were not due to the initial choice of model architecture. This is also demonstrated by Fig. 23, which shows the distribution of filters during training for the shallower UNet when training with the HarP data. It can be seen that, as is evident in the other examples shown, the bottleneck of the model was pruned aggressively early in the pruning, and the first layer at each depth was pruned first – the result shown is for *f* = 8 but the pattern was the same for all values of *f* considered.

**Figure 22:**
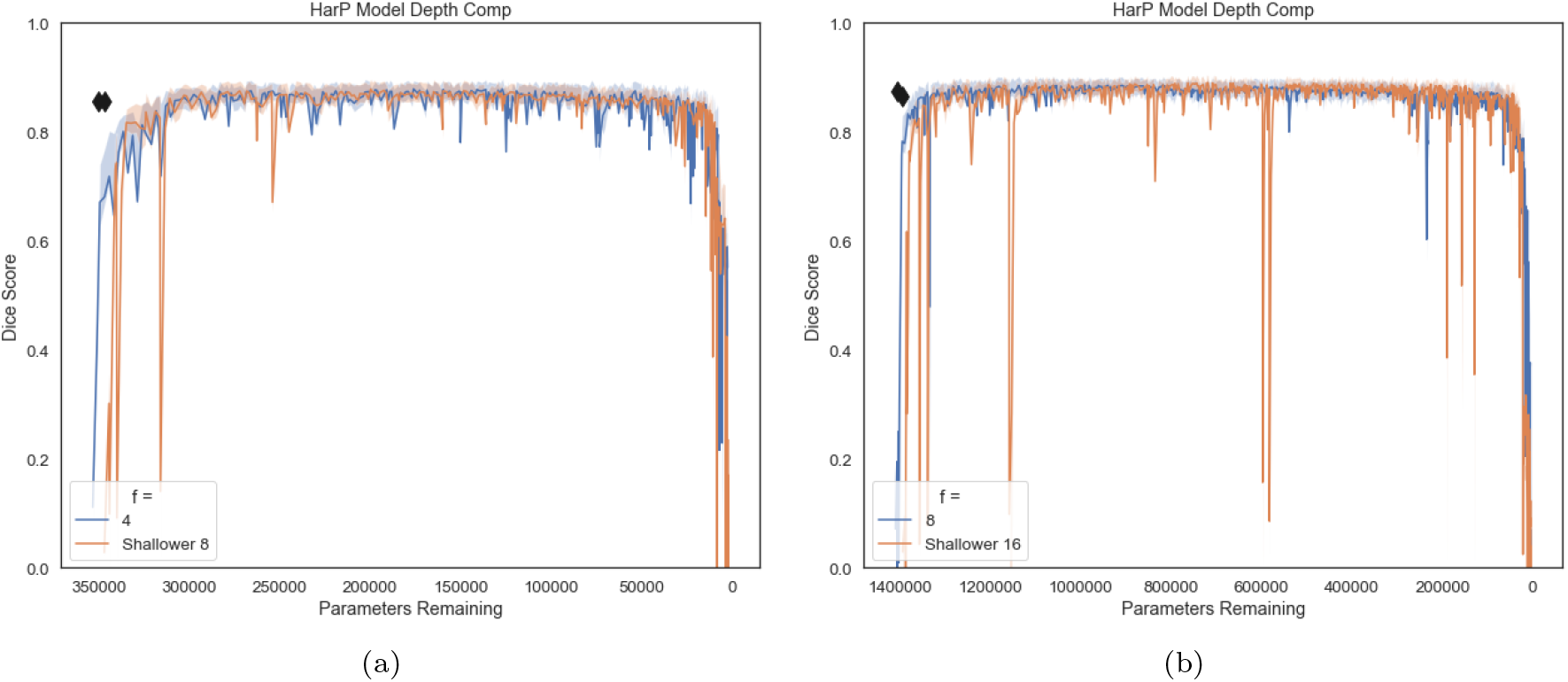
Direct comparison of pruning the original UNet considered – Fig. 3 – and the shallower UNet – Fig. 20 – for the case where the two models have similar numbers of parameters originally: a) compares the UNet with *f* = 4 to the shallower UNet with *f* = 8. b) compares the UNet with *f* = 8 to the shallower UNet with *f* = 16. The black diamonds represent the original *Standard UNets* trained to convergence.

**Figure 23:**
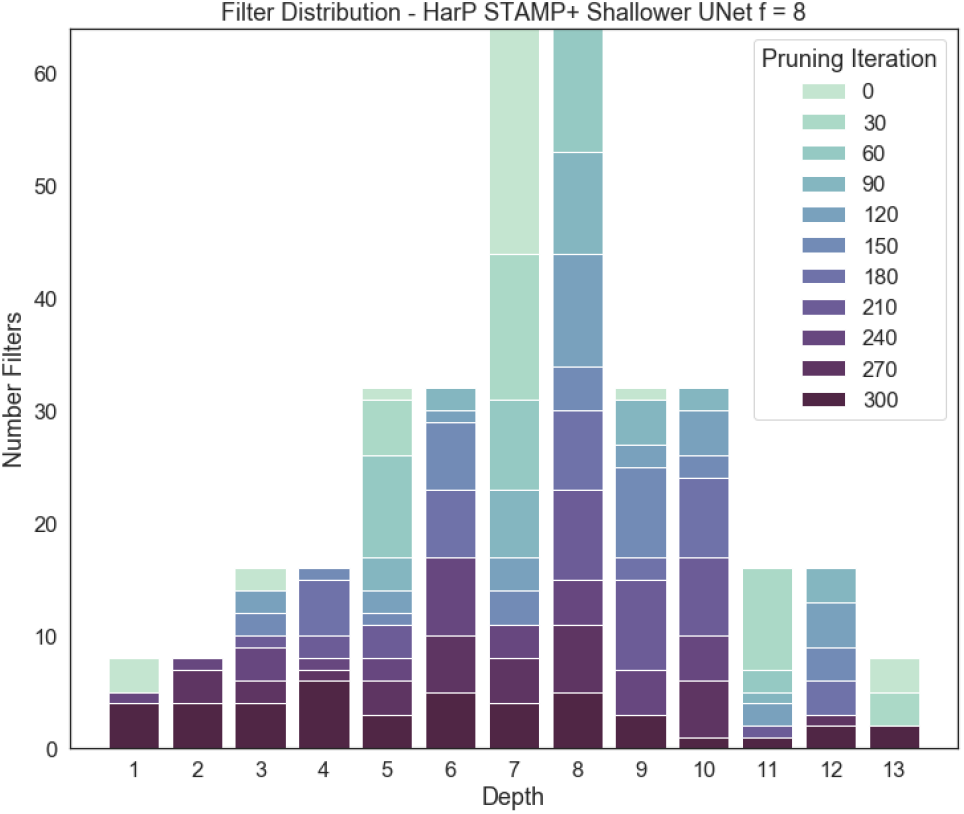
The distribution of the filters with network depth as the shallower model is pruned for hippocampal segmentation on the HarP data, where the darker the shade of the filter block, the longer the filters at that depth were maintained in the model.

### 9.4. Method Comparison

In Fig. 24 the results achieved using different pruning methods are compared: *STAMP+*, *PruneFinetune*, and *STAMP*. First, it can be seen, by comparing *STAMP* with and without targeted dropout, that the targeted dropout both improved the model’s performance (from 0.844 ± 0.03 to 0.879 ± 0.02, *p* < 0.001) and additionally made the training more stable. This shows that the targeted dropout successfully made the model more robust to being pruned, even with this relatively simple task.

**Figure 24:**
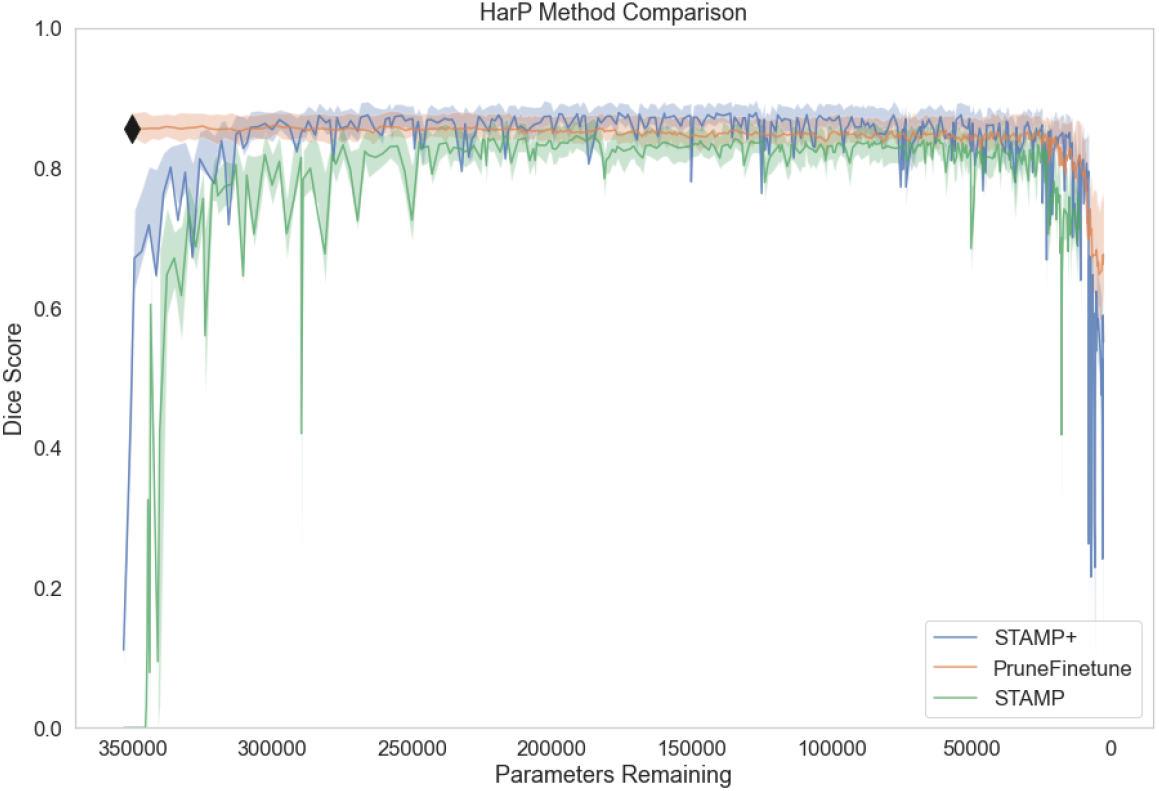
Comparison of pruning the HarP data with *f* = 4 for our proposed method, *STAMP+*, *PruneFinetune* and *STAMP* (i.e. without targeted dropout - green blocks from Fig. 1 removed). The black diamond indicates the performance of the *Standard UNet* trained to convergence without any pruning – *f* = 4.

Comparing *STAMP* to *PruneFinetune*, we can see that pruning the already converged model leads to more stable training than *STAMP* (Fig. 24). However, the removal of a single filter at a time is very conservative compared to the normal approaches taken in the literature, where a percentage of filters, or all filters under a threshold value are removed (Cun et al., 1990), which we would then expect to be less stable. Furthermore, the recovery time between filter prunings is much longer, as the model is allowed to recover entirely before additional pruning is performed (representative training graphs can be found in the supplementary material). Finally, the original had to be trained to convergence before the network could be pruned, and so *PruneFinetune* has a much larger computational cost while performing less well than the proposed method of *STAMP+* (0.860 ± 0.03, 0.879 ± 0.02, *p* < 0.001). Therefore, the proposed method allows better performance to be achieved without having to train the original model.

### 9.5. Training Graphs

Figure 25 shows representative training curves for the two approaches to pruning, with a) showing *STAMP+* during training and b) *PruneFinetune*. The region indicated in red shows the epochs used to train the initial *Standard UNet* model which was used as the initialisation for the pruning process. It can clearly be seen that *STAMP+* requires a fraction of the epochs to train and that the validation performance is much more stable throughout the process.

**Figure 25:**
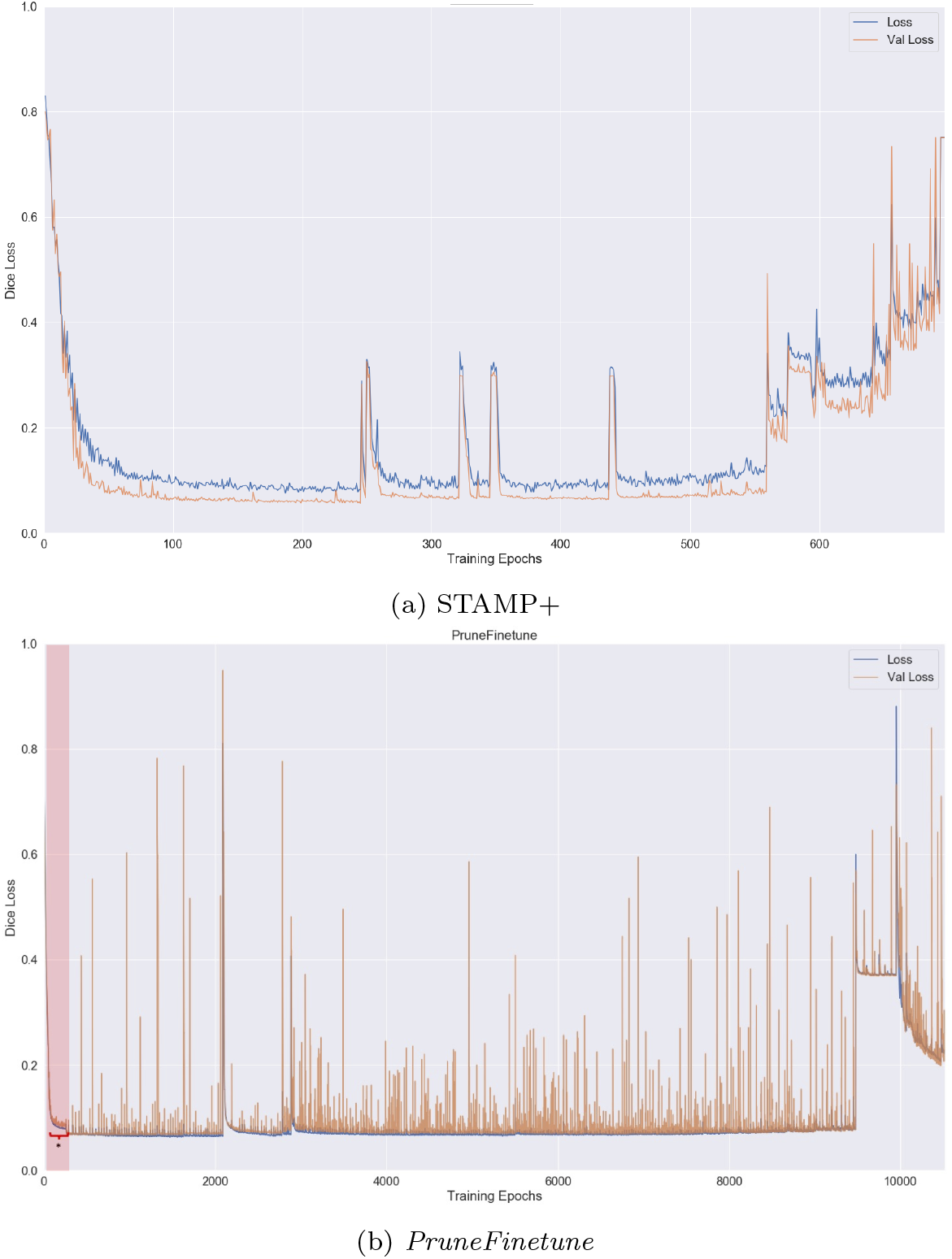
Training and validation loss curves for a) *STAMP+* and b) *PruneFinetune* for the HarP data. The region indicated in red for *PruneFinetune* indicates the epochs for training the initial UNet model.

### 9.6. IXI Subcortical Segmentation

Table 2 shows the results for the IXI subcortical segmentation with increasing amounts of training data, broken down by subcortical region being segmented. It can be seen that when there is insufficient training data to train the *STAMP+* method, the failure mode is the same as the *Standard UNet*, with the network failing to segment one of the subcortical regions.

**Table 2:** Table showing the Dice scores of the different subcortical regions with different amounts of training data for the testing data coming from the Guy’s site, comparing *Standard UNet* and *STAMP+*.

| Number of Training Images | Standard UNet |  |  | STAMP+ |  |  |
| --- | --- | --- | --- | --- | --- | --- |
|  | Caudate | Putamen | Thalamus | Caudate | Putamen | Thalamus |
| 50 | $0.28 \pm 0.07$ | $0.19 \pm 0.03$ | $0.14 \pm 0.04$ | $0.78 \pm 0.04$ | $0.75 \pm 0.04$ | $0.0 \pm 0.0$ |
| 100 | $0.54 \pm 0.12$ | $0.65 \pm 0.08$ | $0.75 \pm 0.06$ | $0.86 \pm 0.02$ | $0.89 \pm 0.04$ | $0.93 \pm 0.02$ |
| 150 | $0.46 \pm 0.06$ | $0.78 \pm 0.06$ | $0.21 \pm 0.07$ | $0.88 \pm 0.02$ | $0.90 \pm 0.03$ | $0.92 \pm 0.02$ |
| 200 | $0.032 \pm 0.01$ | $0.0 \pm 0.0$ | $0.82 \pm 0.02$ | $0.90 \pm 0.02$ | $0.91 \pm 0.03$ | $0.94 \pm 0.01$ |
| 250 | $0.90 \pm 0.03$ | $0.080 \pm 0.05$ | $0.0 \pm 0.0$ | $0.89 \pm 0.02$ | $0.92 \pm 0.03$ | $0.94 \pm 0.01$ |

### 9.7. Evaluation Metric

Dice scores were reported throughout this work. The results for comparing the *Standard UNet* and *STAMP+* were evaluated using different metrics for segmentation performance, and are shown in Fig. 26 for the hippocampal segmentation task utilising the HarP data. It can be seen that the pattern of results was consistent across the choice of evaluation metric.

**Figure 26:**
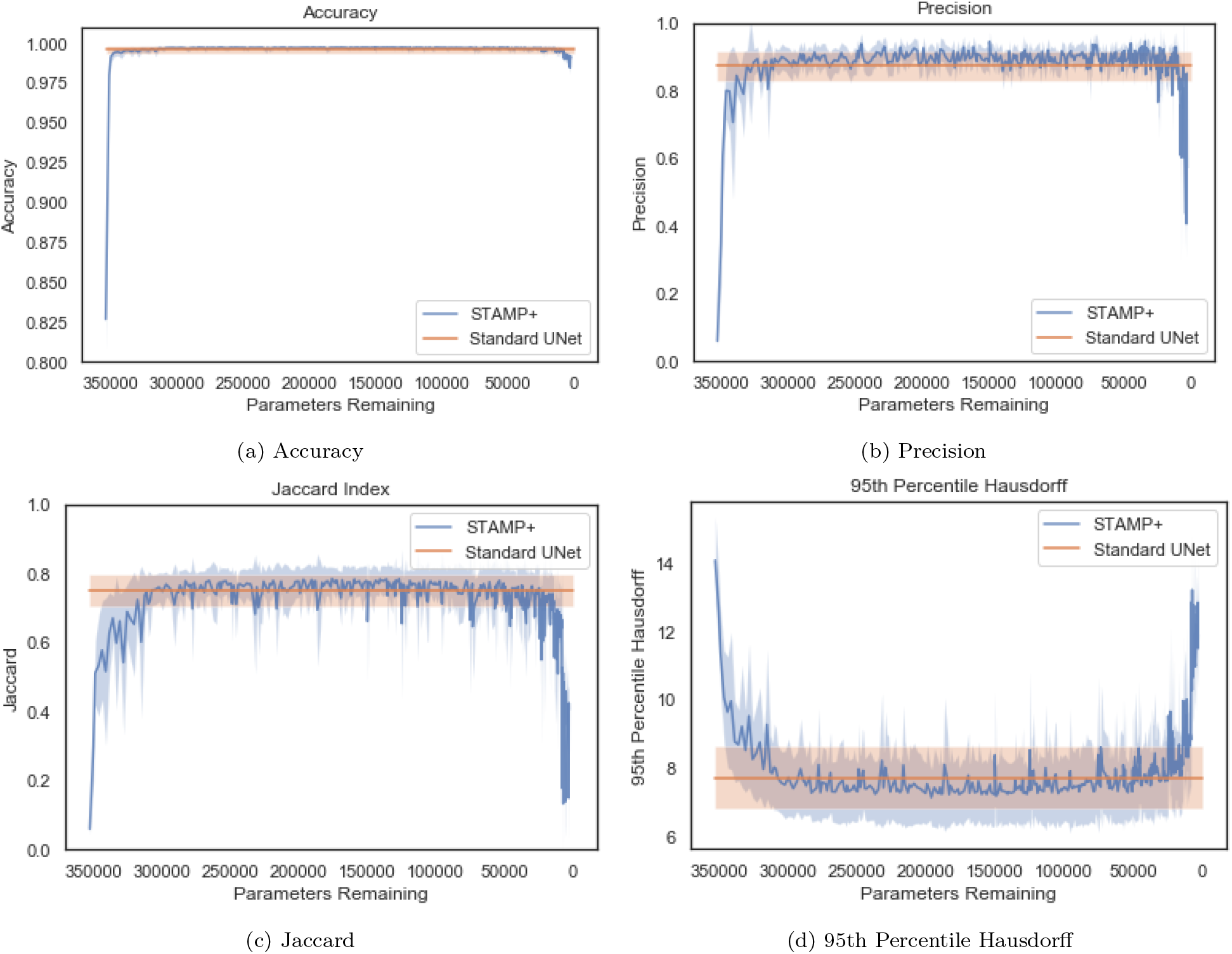
Results with different evaluation metrics, comparing *Standard UNet* and *STAMP+*.

### 9.8. Targeted Dropout Values

Remembering that the UNet architecture conventionally has doubling features with increasing model depth and halving resolution, Fig. 27 shows the adaptive targeted dropout probability values at each layer depth through training, for a model trained with *f* = 8, with every 30 iterations being reported. The *x* value corresponds with the location model, as indicated by the model architecture (Fig. 3). It can be seen that until the model becomes very small (highly pruned, iterations higher than 600) the distribution of the calculated values was very similar between iterations, with a tendency for the bottleneck of the model to have the highest dropout value. This was to be expected, as this is where most filters in the model are located and, thus, where there is likely to be the greatest level of redundancy. It can also be seen that towards the end of training the dropout probabilities for some of the depths are 0. These corresponded to the depth where only a single filter remains, thus to prune (or to apply dropout) would break the model and information would be unable to flow.

**Figure 27:**
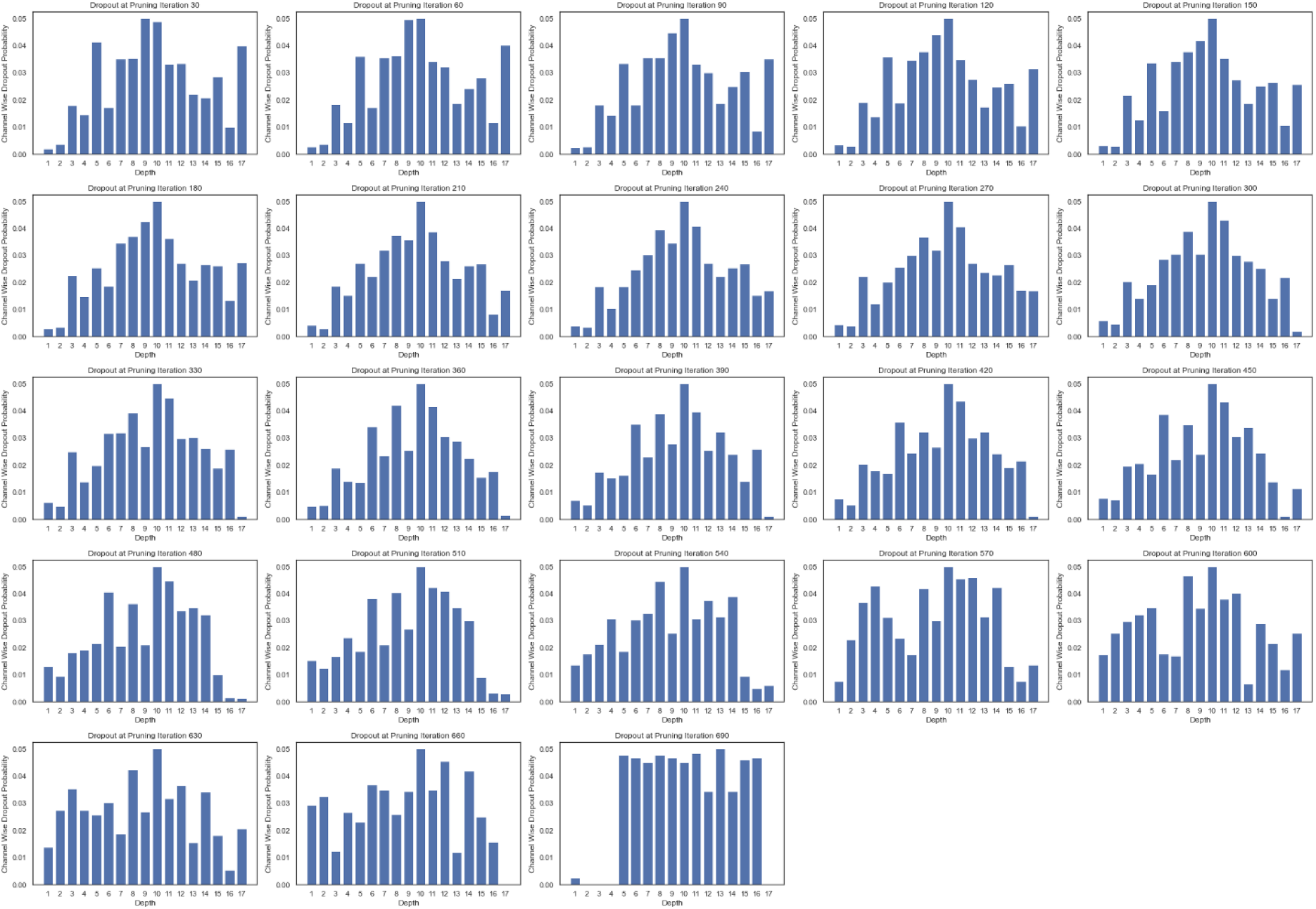
The adaptive targeted dropout probability values at each layer (depth) for every 30 epochs of training on the HarP data – *b_drop_* = 0.05 – for *f* = 8. It can be seen that the dropout values are quite consistent across pruning iterations until the network becomes very small in size. When a layer has a single filter at that depth, the dropout value becomes 0, guaranteeing that information can be propagated through the model.

### 9.9. Filter Survival

Figure 28a shows the distribution of the number of filters at each depth as the network was pruned. The darker the section of the bar, representing the number of filters at a given layer, the longer the filters remained in the model architecture. As would be expected, as the pruning was based on the activation magnitude values, the filters in the bottleneck of the network are pruned aggressively first, corresponding to the high dropout values. It can also be seen across the network that the first layer within a pair of layers, at a given depth, is pruned more quickly than the second, for both the encoder and the decoder.

**Figure 28:**
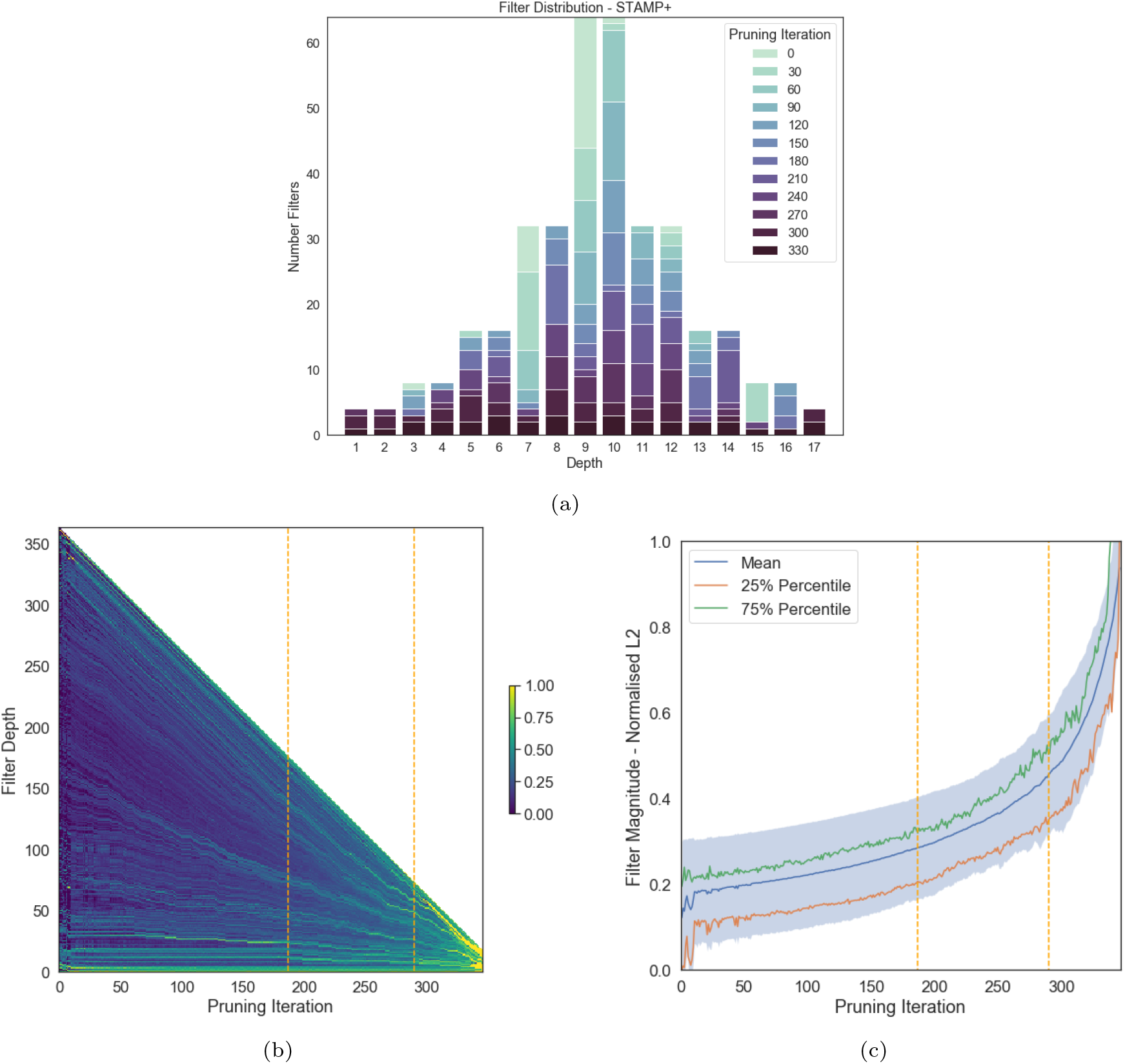
a) Shows the distribution of the filters with network depth as the model is pruned, meaning that the darker the shade of the filter block, the longer the filters at that depth were maintained in the models. b) Shows the magnitudes of the activations, averaged across the training data, maintained in the model as it was pruned, where filter depth is the count of filters from the input. It can be seen that the lower magnitude activations were pruned first and the average value of the activation increased as the model was pruned. The first vertical dashed line at 182 filters corresponds to the distribution of filters which gave the best performance on the testing data. The second dashed line corresponds to the distribution of the filters for the smallest model that was able to complete the segmentation successfully, with no substantial difference in performance from the best performing model. c) Shows the average filter magnitude, and the lower and upper quartile bounds with pruning iteration. It can be seen that the average value increased consistently with pruning iteration.

**Figure 29:**
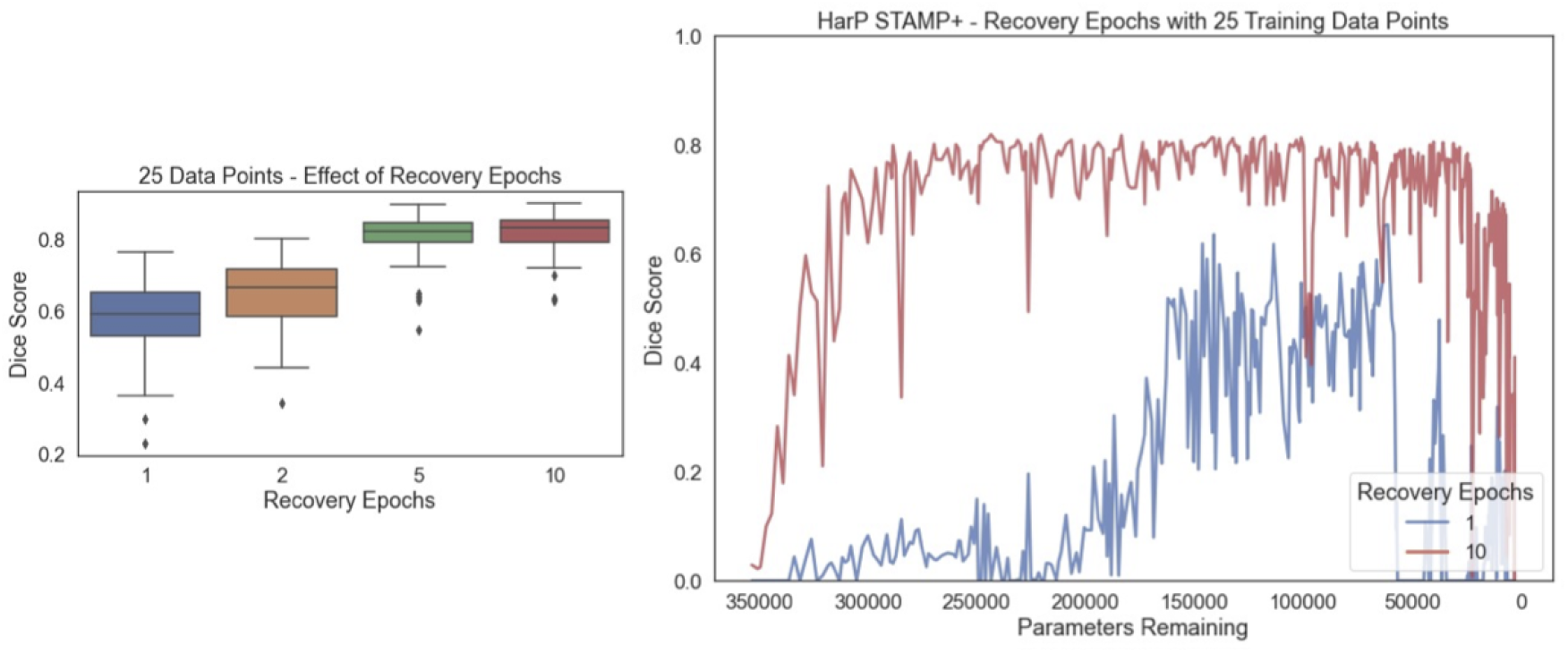
Effect of increasing the number of recovery epochs on the segmentation performance on the HarP data with 25 training examples. The box plot shows the best performance achieved, chosen on the validation data, and the lineplot shows the pruning dice score for 1 and 10 recovery epochs as the training progressed, showing the mean value as the number of parameters decreased.

**Figure 30:**
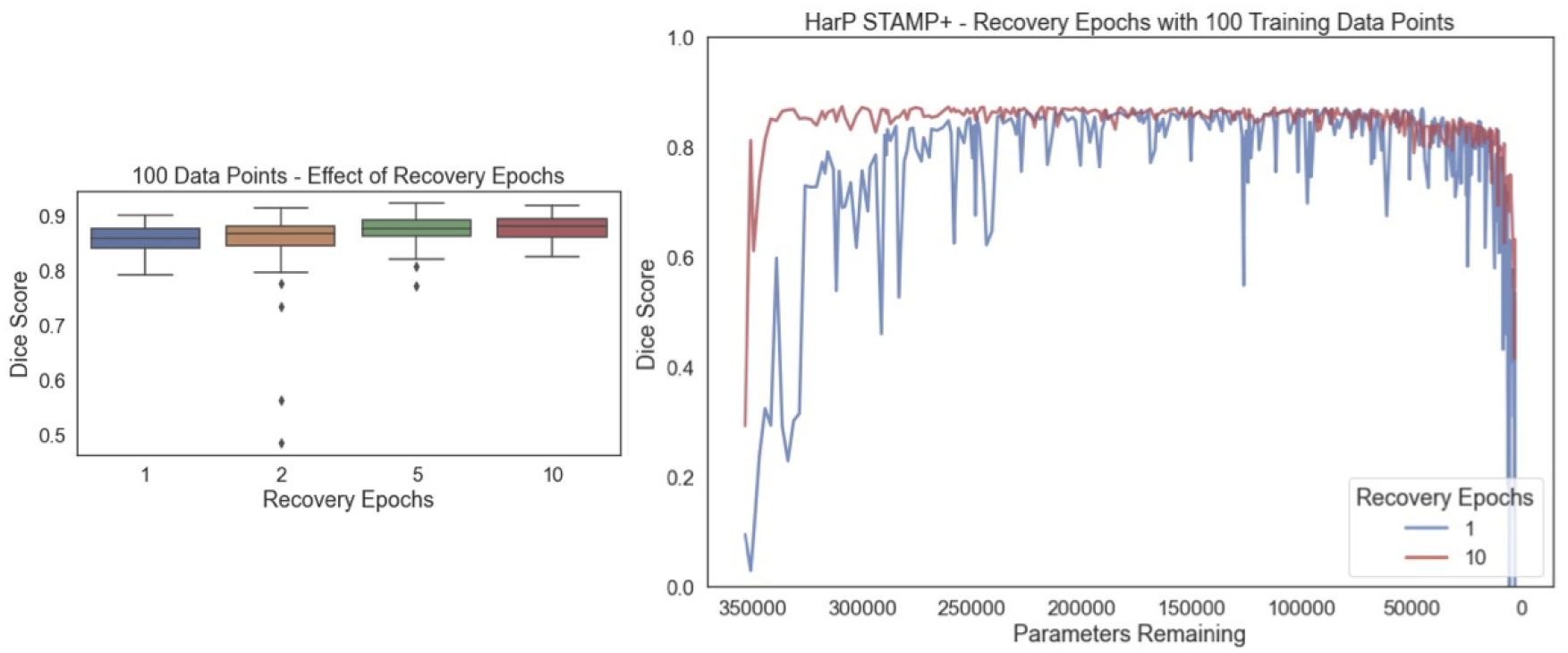
Effect of increasing the number of recovery epochs on the segmentation performance on the HarP data with 100 training examples. The box plot shows the best performance achieved, chosen on the validation data, and the lineplot shows the pruning dice score for 1 and 10 recovery epochs as the training progressed, showing the mean value as the number of parameters decreased.

Figure 28b shows the magnitudes of the filters: 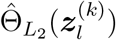 averaged across the testing dataset as the model is gradually pruned. It can be seen that as the model is gradually pruned, the average value of the remaining filters increased and the high magnitude filters remained throughout the pruning. This can be seen by considering the strong bright lines which are maintained throughout the pruning: given that a filter was initially important, it remained important throughout the training and the activations only increased in value.

The first dashed vertical line at iteration 182 represents the point in the pruning where the model had the best performance on the testing data (49% of the filters of the original model). It can be seen that the majority of the low-magnitude filters (assumed to be uninformative) have been removed, showing that more than half of the filters in the model could be removed without having a negative impact on the segmentation. The second dashed line represents the smallest network (20% of the filters of the original model) for which the performance was not significantly worse than the best performing model. This indicates that the network can be reduced to a small number of filters, where all of the activations were playing a more important role in producing the outputs.

These results clearly demonstrate that the proposed method could be utilised to simultaneously train and prune a UNet, while working in a low-data regime, as is common in medical imaging. The results also indicate the benefit of the addition of the targeted dropout, showing an increased performance, even on this relatively easy task.

### 9.10. Recovery epochs

We repeated the experiment presented in Section 3.4.1 on 25 and 100 data points respectively. It can be seen that the pattern is the same as that presented for 50. The more data points available for training, the lower the impact of the number of recovery epochs became.

## Footnotes

1 Data downloaded from: https://brain-development.org/ixi-dataset

2 For details see: https://fsl.fmrib.ox.ac.uk/fsl/fslwiki/fsl_anat

## Notes

### Competing Interest Statement

The authors have declared no competing interest.

### Summary of Updates

figures updated

